# Trade-off between resource excretion and reliance maintains biodiversity of a cross-feeding mutualism

**DOI:** 10.64898/2026.11.24.768994

**Authors:** Yihe Yu, Ave T. Bisesi, Arjun Mandyam Dhati, Iona Kambouridis, Hiya Parikh, Akshit Goyal, Clare I. Abreu

## Abstract

Microbial mutualisms, in which species exchange resources through cross-feeding, are common in nature. Yet the stability of mutualistic interactions is debated, and cross-feeding communities are vulnerable to cheaters that benefit from the excretions of others but reduce or cease their own excretion of resources, which can lead to community collapse. Here, we demonstrate with a mathematical model that in addition to decreased excretion, increased reliance on cross-fed resources can also lead to community collapse. In a two-species model of bacteria that metabolizes lactose and provides yeast with galactose, in exchange for amino acids from yeast, we show that the set of excretion and reliance parameters that maximizes biomass is adjacent to a region of community collapse in the phase diagram. By expanding the model to include multiple bacterial auxotrophs competing for lactose and either separate or overlapping amino acids, we also demonstrate that a trade-off between reliance and excretion can enable coexistence within the bacterial mutualist guild. Finally, we show that intraguild coexistence of bacteria favors yeast, increasing biodiversity but decreasing total community biomass. Our results argue that the question of cheaters should be paired with analysis of reliance on mutualistic exchange, and that both of these parameters modulate biodiversity and the stability of mutualisms to collapse.

## INTRODUCTION

Most microbes live in diverse communities where environmental conditions, including resource availability, can undergo significant fluctuations [Abreu et al. 2020]. Despite the fact that resource availability can be unpredictable, 26% to 46% of bacterial genotypes are auxotrophic, or unable to produce all required amino acids [Yu et al. 2022], and thus can survive only in environments where amino acids are supplied at sufficiently high concentrations. While these resources may often be obtained from the abiotic environment [Hoek et al. 2016, Hammarlund et al. 2019], microbial cross-feeding of both essential and non-essential resources, including acids, sugars and vitamins, is also common [Kost et al. 2023]. This type of metabolic cross-feeding is typically the result of strains metabolizing externally available resources and leaking (i.e. excreting) waste or overflow products to maintain intracellular homeostasis [McKinlay 2023]; these byproducts can then be used by other strains. For example, some bacteria excrete galactose during lactose metabolism (**Fig. 1A**) [Ponomarova et al. 2017, Gabrielli et al. 2023], while for others carbon conversion during growth on sugars can lead to the excretion of organic acids [Goldford et al. 2018, Amarnath et al. 2023, Paczia et al. 2012] and nitrogen overflow processes can drive the externalization of excess amino acids in yeast [Ponomarova et al. 2017, Paczia et al. 2012, Correia-Melo et al. 2023]. The strength and stability of positive (e.g. mutualistic) interactions between microbial strains therefore generally depends on the extent to which strains receive and rely on key resources from partners versus the abiotic environment.

**Figure 1.**
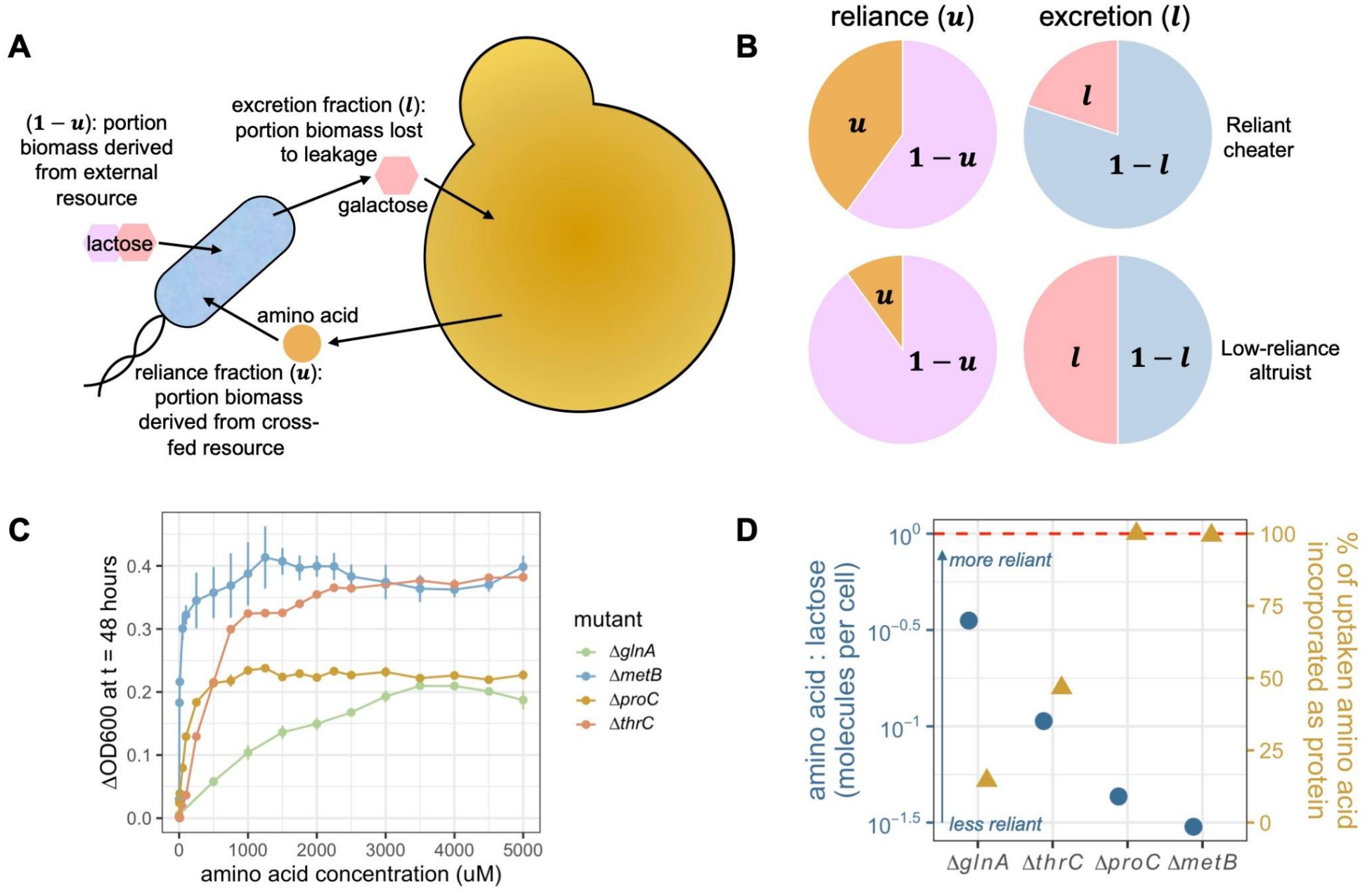
Proteinogenic demands and biosynthetic pathways shape relative auxotroph reliance on externally supplied amino acids. **A:** Schematic of a model probiotic system of bacteria and yeast, where bacteria metabolize lactose and excrete galactose, making it available to yeast, which cross-feed amino acids required by the bacteria. **B**: Examples show different combinations of reliance and excretion fractions. A “reliant cheater” has a high reliance on cross-fed amino acids relative to lactose and excretes less galactose. A “low-reliance altruist” relies less on amino acids and excretes more galactose. **C**: The maximum OD600 of different *E. coli* single amino acid auxotrophs when grown in a lactose-based media with varying concentrations of their required amino acid externally supplied. Error bars indicate standard error of the mean across three replicates. **D:** *Left axis*: The log-transformed ratio of amino acid molecules to lactose molecules required to make one *E. coli* cell as predicted by flux balance analysis (FBA) for four single amino acid auxotrophs, where values below 1 indicate that the cell requires more lactose than amino acid. *Right axis*: The percentage of imported amino acid incorporated into protein rather than used as a precursor for other biosynthetic pathways, as predicted by FBA.

While ecological theory has traditionally focused on predicting the dynamics of systems dominated by competition, recent work has increasingly examined how mutualistic interactions like metabolic cross-feeding can promote diversity and stability in microbial communities [Chacón et al. 2021, Goldford et al. 2018, Goyal et al. 2018]. For example, both theory and experiments have demonstrated that cross-feeding via leakage is sufficient to support diverse communities even during growth on a single external resource [Goldford et al. 2018, Goyal et al. 2018], while other models have suggested that diversity can be maximized when external resources are supplied at high rates and microbes excrete a large fraction of the consumed resource [Marsland et al. 2019]. In these frameworks, cross-feeding is largely facilitative, where excreted metabolites are shared but no species strictly depends on another for growth. Yet sufficiently high excretion rates can also stabilize more stringent forms of mutualism such as obligate cross-feeding mutualisms, which are otherwise susceptible to collapse because neither partner can survive alone [Shou et al. 2007]. However, while it is clear that excretion is a central determinant of coexistence for positively interacting microbes, many of these models have omitted the fact that a strain’s reliance on cross-fed metabolites relative to externally-supplied resources will depend on both genetic and environmental context [Baichman-Kass et al. 2026]. The stability of mutualism is often evaluated in the face of cheating, where a strain benefits by reducing its production of shared resources without changing its dependence on its partner [Estrela et al. 2016]. The inclusion of reliance represents the alternative possibility that mutualism breakdown may proceed through reduced dependence on the shared resource (**Fig. 1B**). While reduced reliance does not necessarily preclude the coexistence of obligately cross-feeding partners, in the extreme case, freeing only one partner from dependence on an obligate mutualism, for example by providing the cross-fed resource externally, can drive system collapse [Hammarlund et al. 2019]. Both excretion and reliance thus represent distinct interaction dimensions that should determine the stability of obligate cross-feeding systems.

Because neither species can persist alone, the outcome of pairwise cross-feeding under obligate conditions has only two possible qualitative outcomes: coexistence or collapse. When coexistence is possible, a range of system- and model-specific details may dictate quantitative outcomes like the total yield and relative abundance of interacting strains. However, while most studies of mutualism have focused on pairwise interactions [Chacon et al. 2021], multi-member mutualist guilds are more common in nature [Stanton 2003]. Despite the fact that members of an obligate mutualist guild will necessarily compete for essential resources and can have both overlapping and non-overlapping niches [Johnson and Bronstein 2019], intraguild diversity is still common for these interactions, in part because more diverse mutualistic communities are buffered against collapse by functional redundancy [Vidal et al. 2020]. Existing theory suggests that a range of trade-off types can promote intraguild coexistence for obligate interactions, such as dependence on different cross-fed resources from a shared, growth-limiting partner [Hammarlund et al. 2021] or growth rates differences across external and cross-fed resources [Johnson and Bronstein 2019]. Classic resource competition models that do not include cross-feeding show that a “gleaner-opportunist” trade-off [Grover 1990] between growth in high and low resource concentrations allows competing species to coexist and thus may be relevant for the maintenance of intraguild diversity. However, since obligate cross-feeding limits overall growth to match a partner [Chacon et al. 2021], growth trade-offs within a guild may generally be less relevant under these mutualistic conditions than they are during resource competition between strains. Instead, we expect trade-offs between excretion and reliance to determine qualitative and quantitative outcomes in obligate systems, both between pairwise mutualists and within mutualist guilds.

Here, we consider the importance of excretion and reliance in an obligate cross-feeding mutualism where yeast excrete amino acids that are consumed by bacteria and bacteria excrete galactose that is consumed by yeast (**Fig. 1A**), an interaction relevant to environments where lactose is provided as an external resource [Ponomarova et al. 2017, Blasche et al. 2021, Gabrielli et al. 2023]. First, we experimentally and computationally demonstrate variability in reliance on different amino acids across a library of bacterial auxotrophs (**Fig. 1C-D**). We then develop a consumer-resource model that incorporates both excretion and reliance. These two parameters alone determine the analytical solution for equilibrium biomass of pairwise cross-feeding; surprisingly, biomass is maximized in the region of parameter space adjacent to a region of system collapse. Extending the model to include multiple bacterial auxotrophs that cross-feed with yeast, we find that trade-offs between excretion and reliance enable greater coexistence among auxotrophs during cross-feeding than when amino acids are supplied externally. In both pairwise and multi-auxotroph cases, the excretion-reliance trade-off determines both equilibrium biomass and biodiversity among bacteria and yeast. While a gleaner-opportunist trade-off may modulate coexisting frequency among bacterial auxotrophs, we observe that it does not influence total system or yeast biomass. The excretion-reliance trade-off, on the other hand, modulates biomass such that coexistence among competing bacterial auxotrophs increases the biomass of the shared yeast partner with which they cross-feed. Our results demonstrate that reliance on cross-fed relative to external resources is an important parameter that modulates community outcomes, and which should be included in future models of cross-feeding in particular, and of multiple essential resources more generally.

## RESULTS

Inspired by the importance of bacterial-yeast cross-feeding interactions across a range of microbiomes [Tamang et al. 2016, Menezes et al. 2018, Blasche et al. 2021] (**Fig. 1A**), we first generated a library of bacterial amino acid auxotrophs with the potential to cross-feed with yeast isolates. We performed a series of titration experiments to identify the concentrations of respective amino acids required to support carbon-limited growth of each auxotroph (**Methods**). We observe significant differences in maximum possible yield and limiting amino acid concentrations between auxotrophs; for example, very little methionine (<250 uM) is necessary to support growth of the Δ*metB* auxotroph, while roughly ten times as much glutamine is required for robust growth of the Δ*glnA* auxotroph (**Fig. 1C**). Using flux balance analysis (FBA) (**Methods**), we investigated the metabolic bases of these differences by calculating the predicted intracellular demand for each amino acid relative to lactose (**Fig. 1D**). FBA predicts that the Δ*glnA* mutant requires twice as much glutamine as lactose to support its growth due to the importance of glutamine as a biosynthetic precursor (**Fig. 1D**), while only 1/30th the amount of methionine is necessary for the Δ*metB* mutant because nearly all imported methionine is assimilated into protein, with the proteome therefore setting the level of methionine demand. In line with previous work [Baichman-Kass et al. 2026, Hammarlund et al. 2021, Hammarlund et al. 2019], these results emphasize that the identity of cross-fed nutrients should have significant consequences for the success of mutualistic interactions. Given the high levels of predicted auxotrophic reliance, higher levels of excretion of a biosynthetically costly compound like glutamine might be required from a mutualistic partner to stabilize obligate cross-feeding interactions. We therefore sought to examine the relative importance of excretion and reliance for mutualistic stability using a modified consumer-resource model.

### Mutualistic reliance and metabolite excretion shape the stability of mutualistic cross-feeding in a two-species consumer-resource model

To predict how resource reliance and metabolite excretion would impact the stability of obligately mutualistic cross-feeding, we began with a two-species, three-resource consumer-resource model (**Methods**). In the model, bacteria and yeast trade galactose andamino acid in a minimal lactose medium; under these conditions, the strains engage in obligate mutualism, and one cannot grow without the other (**Fig. 2A**). Based on existing theory on the stability of obligately mutualistic systems [Shou et al. 2007], we expected that both high levels of reliance and excretion should be required to support robust growth and coexistence in our system. Given that previous microbial cross-feeding models have explicitly considered metabolite excretion but not reliance, we first sought to understand how our reliance parameter *u* influenced mutualistic stability. In the model, both lactose and amino acid are required for bacterial growth; *u* is the fraction of bacterial biomass derived from amino acid supplied by yeast, while 1 − *u* is the fraction derived from external lactose.

**Figure 2.**
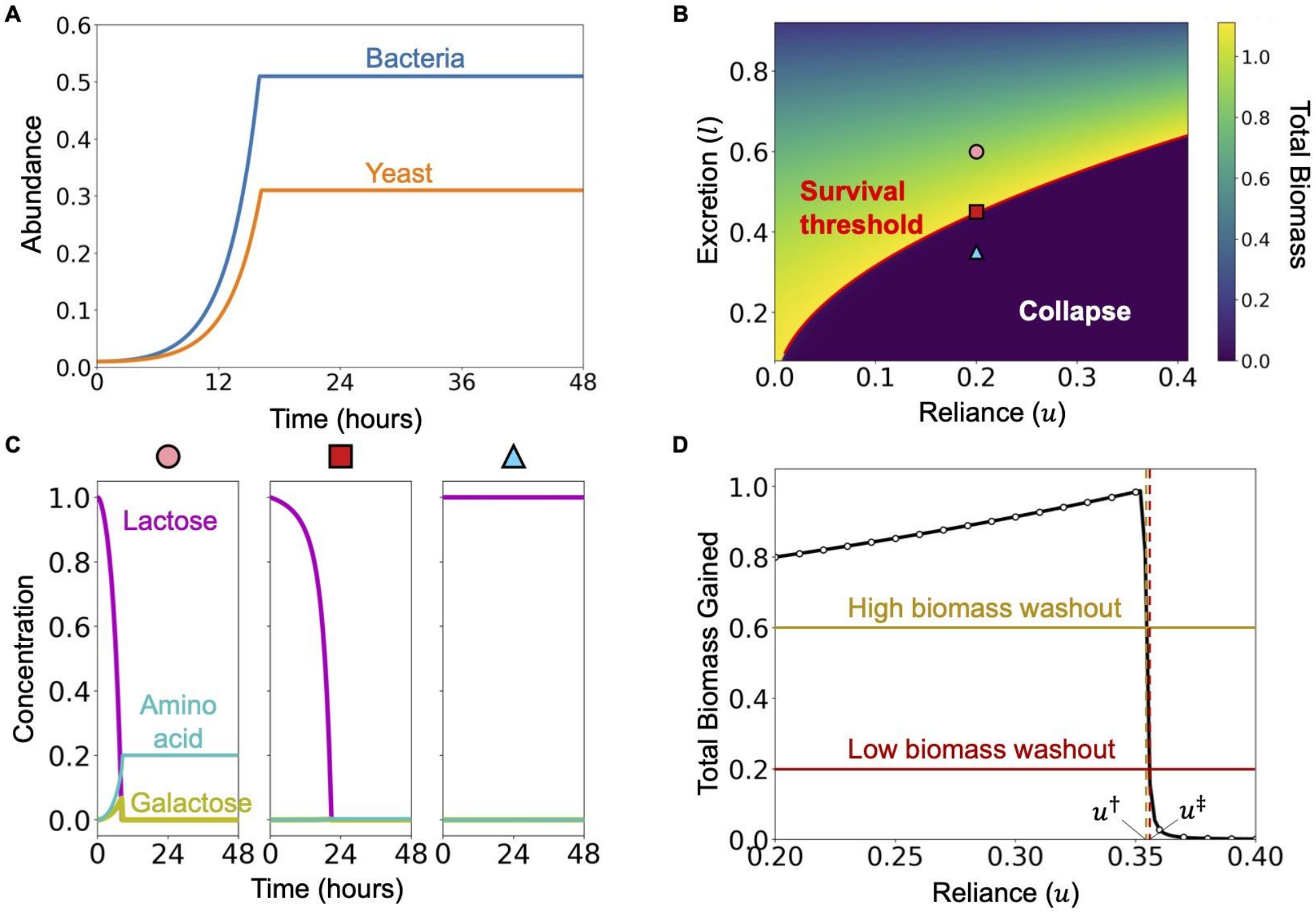
Mutualistic reliance sets the lower limit on excretion of cross-fed resources. **A**: Simulated time-series dynamics of yeast and bacterial biomass in a batch culture consumer-resource model when lactose is supplied as the sole carbon source. **B**: Total microbial biomass across varying bacterial reliance and excretion. The red boundary delineates the analytical excretion-reliance threshold below which co-culture growth is impossible. **C:** Time-series dynamics of three resources in a batch culture consumer resource-model at different levels of bacterial excretion (reliance held constant). **D**: Total microbial biomass across varying bacterial reliance. The dashed vertical lines correspond to the reliance threshold above which co-culture growth is impossible. Solid horizontal lines correspond to the amount of total biomass required to offset varying levels of system perturbation (i.e. dilution rate).

We solved the model analytically to identify the minimum level of reliance *u*\* required to enable co-culture growth (**Methods**). We found that *u*\* depends primarily on the leakage parameters *l*, which represent the fraction of consumed resources excreted by both bacteria and yeast, and on the initial concentrations of the two cross-fed resources. Growth in our model can only be considered obligately mutualistic when the initial concentrations of both cross-fed resources are at or near zero, reducing our analytical solution for *u*\* to a relationship between reliance and excretion without dependence on any other parameters (**Methods**). We extended this analytical solution to quantify total system biomass at steady state given a *u* value above or below *u*\*, finding that biomass can likewise be predicted solely by reliance and excretion parameters, along with a handful of other parameters assumed to be constant in our system (**Methods**). By solving for total biomass across a range of *l* and *u* values, we observe a region of system collapse predicted by *u*\* (**Fig. 2B**), such that increasing reliance increases the amount of excretion required to support co-culture growth. While this is an intuitive finding, we were surprised that our analytical results further suggest that system biomass is maximized in the region of reliance-excretion parameter space nearest the collapse boundary. Above this boundary, biomass decreases as excretion increases and/or reliance decreases (**Fig. 2B**). We also observe that bacterial frequency increases as excretion decreases, reaching highest frequencies at the collapse boundary (**Fig. S1**).

To understand why small changes in excretion and reliance parameters shift our mutualistic system from optimal biomass to collapse, we investigated the resource dynamics in these different regimes (**Fig. 2C**). The region of system collapse is defined by the inability of the cross-feeding strains to deplete externally supplied lactose; simulations verify that whenever growth is possible, lactose is exhausted (**Fig. 2C**, circle and square). The trend for the final concentration of amino acid is inverted: when growth is impossible, this is because the rate of amino acid excretion by yeast is insufficient to support the growth of bacteria and the amino acid thus serves as the limiting reagent for the system. By reducing the efficiency with which lactose can be converted into bacterial biomass, high levels of bacterial excretion increase the amount of excess amino acid present in the media at the end of growth (**Fig. 2C**, circle). As such, the collapse boundary functionally demarcates the transition of the cross-feeding system from being amino acid-to lactose-limited. Biomass is maximized directly above this boundary because in this region both resources are depleted similarly, optimizing total resource usage (**Fig. 2C**, square). Given that high excretion levels reduce biomass production, we wondered how sensitive the *u*\* boundary would be to biomass washout from dilution.

We therefore investigated the consequences of varying biomass washout levels on the *u*\* collapse boundary by plotting our analytical solution for total biomass across increasing reliance *u*, with excretion held constant (**Fig. 2D**). As expected, we find that collapse occurs above a given value *u*\* and that total biomass is maximized when reliance is slightly less than that threshold value. The sharp drop-off of this biomass curve clarifies the relative importance of the dilution factor. By considering two possible (but arbitrary) washout levels - a low perturbation state, where total biomass need only reach a value of 0.2 each cycle to prevent washout from dilution, and a high perturbation state, where total biomass would need to reach a value of 0.6 to offset dilution rate - we observe that the levels of mutualistic reliance *u*^†^ and *u*^‡^required to support growth at these baseline biomasses are nearly identical (**Fig. 2D**). Therefore, our model suggests that the exact rate of dilution should be of little consequence to the threshold level of mutualistic reliance or the overall stability of the mutualistic system. Together, these results demonstrate that excretion and reliance are the fundamental parameters tuning the growth and stability of our cross-feeding model across a wide range of parameter space, supporting the importance of their inclusion for predicting mutualistic dynamics.

### Cross-feeding supports greater coexistence between competing niche-separated auxotrophs

Given the observed importance of excretion and reliance parameters in our two-species, three-resource system, we next sought to understand how the inclusion of these biological details would shape the stable coexistence of multiple competing bacterial strains. Since previous work [Hammarlund et al. 2021] has demonstrated that coexistence is possible when bacteria require different amino acids from their mutualistic partner (i.e. niche separation), but that coexistence is more challenging when amino acids are externally supplied, we amended our model in two ways. First, we expanded it to include an additional amino acid and the auxotroph requiring it (**Fig. 3A-B, Methods S1**). Under cross-feeding conditions, lactose is again supplied in the medium as the sole initial resource and consumed by two bacterial auxotrophs with different amino acid requirements, threonine or methionine. Yeast are assumed to provide these two amino acids at equivalent rates, proportional to total yeast excretion, to the competing bacterial strains (**Fig. 3A**). As has been shown, we expected that coexistence between competing auxotrophs with amino acid niche separation should be possible across a wide range of parameter space during cross-feeding. In contrast, we anticipated that competitive exclusion should be more common when amino acids are externally supplied at equivalent initial conditions and yeast are absent from the system.

**Figure 3.**
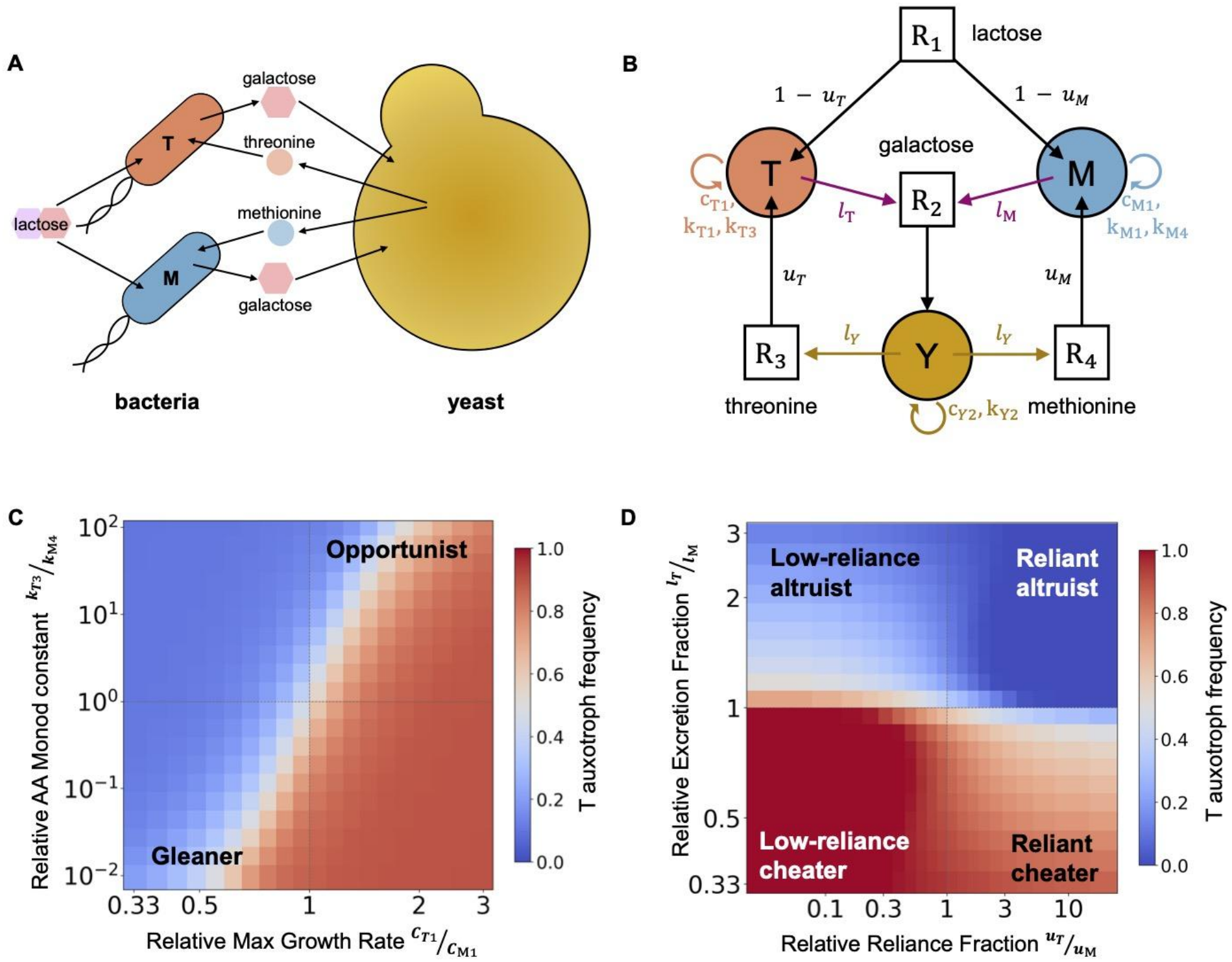
Gleaner-opportunist and reliance-excretion trade-offs predict coexistence in a partitioned mutualist guild. **A-B**: Schematic representation of the 3 strain system in Eqs. S13-S19. Threonine and methionine bacterial auxotrophs compete for lactose but consume separate cross-fed amino acids. **C:** Heat map of the fraction of the bacterial population composed of the threonine auxotroph (T) during co-culture growth as maximum uptake rates and amino acid (AA) half-saturation constants vary. Relative maximum uptake values greater than 1 indicate that T grows faster; relative amino acid half-saturation constants greater than 1 indicate that T requires a higher concentration of its amino acid to reach its maximum growth rate. **D:** Heat map of the fraction of the bacterial population composed of the threonine auxotroph (T) during co-culture growth as excretion and reliance vary. Relative reliance values greater than 1 indicate that T requires more units of amino acid per unit of lactose to generate a single cell; relative excretion values greater than 1 indicate that T excretes a greater percentage of lactose as galactose.

To verify this intuition, we built upon the results of Hammarlund et al. 2021 by investigating the gleaner-opportunist trade-off, usually described in the context of competition for an external resource [Grover 1990, Fridman et al. 2022], under both cross-feeding and non-cross-feeding conditions. In this framework, coexistence between species competing for a shared limiting resource is enabled by differences in growth rates and resource affinities. A strain that can grow quickly at high resource concentrations (“opportunist”) can coexist with a strain that grows more slowly but has higher affinity for the shared resource (“gleaner”). We implemented this trade-off by changing the relative growth rates and amino acid Monod constants of the two auxotrophs (**Fig. 3C, S2**), initializing both the maximum uptake rate and Monod constant at our default parameter values (**Table 1**) and then shifting the trait pair values in opposite directions. For example, for uptake, we varied *c*_*T*_ and *c*_*M*_ such that *c*_*T*_ = 0. 8 + *x* where *c*_*M*_ = 0. 8 − *x* and vice versa. Relative values were set using the threonine auxotroph (T) as the reference, where a relative growth rate or Monod constant greater than one indicates that T has the larger value, while all other parameters are held constant.

**Table 1.** Default values and biological meaning of modeling parameters.

| **Parameter** | **Meaning** | **Default value and unit** |
|---|---|---|
| *N*<sub>*i*</sub> | biomass or abundance of species *i* | initial concentration in simulations: 0.01 biomass for each species present |
| *R*<sub>*j*</sub> | concentration of resource *j* | initial concentrations:<br>lactose (*R*<sub>1</sub>): 1<br>galactose (*R*<sub>2</sub>): 0<br>amino acids (*R*<sub>3</sub>, *R*<sub>4</sub>, *R*<sub>5</sub>): 1e-4 |
| *c*<sub>*ij*</sub> | maximum growth rate of species *i* on resource *j* | 0.8 resource per unit biomass per hour |
| *k*<sub>*ij*</sub> | Monod constant | 0.001 resource |
| *Y*<sub>*ij*</sub> | yield of species *i* on resource *j* | 1 biomass per unit resource |
| *l*<sub>*ij*</sub> | leakage/excretion fraction, or fraction biomass lost to excretion | 0.6 (unitless) |
| *u*<sub>*j*</sub> | reliance fraction, or fraction biomass derived from cross-fed amino acid *R*<sub>*j*</sub><br>(fraction derived from external lactose is 1 − *u*<sub>*j*</sub>) | 0.2 (unitless) |
| *p*<sub>*j*</sub> | the partitioned fraction of amino acid *R*<sub>*j*</sub> yeast makes per unit of its excreted flux.<br>∑<sub>*j*</sub>*p*<sub>*j*</sub> = 1 | $\frac{1}{number\ of\ bacteria\ strains}$ |
| *a* | conversion ratio by *G* of threonine to glycine | 0.5 (units threonine that 1 unit glycine is equivalent to) |

As expected, we find that coexistence is robust during cross-feeding and that which strain dominates is well-predicted by the gleaner-opportunist trade-off boundary demarcating 50/50 coexistence (**Fig. 3C**). This trade-off is not relevant to biomass, however: bacterial auxotrophs change frequencies, but total bacterial biomass and total system biomass remains constant across the tested parameter space (**Fig. S3**). In contrast, when amino acids are externally supplied, competitive exclusion of one of the auxotrophs is the norm, with only narrow regions of coexistence where growth rates are nearly identical and Monod constants are as different as possible (**Fig. S2**). When lactose resource affinity is varied and amino acid constants are fixed, coexistence and exclusion dynamics instead depend only on relative growth rate (**Fig. S4**). This is not surprising given that lactose is always supplied in excess and limiting for only brief portions of growth regardless of whether amino acids are cross-fed or externally supplied, minimizing the relevance of high lactose affinity under both environmental conditions. Furthermore, these results demonstrate that the gleaner-opportunist trade-off can modulate coexistence frequencies of niche-separated auxotrophs cross-feeding with a shared partner, but despite its utility in understanding intraguild dynamics, the trade-off has no effect on the potential for interguild coexistence or collapse of bacteria and yeast.

We next explored whether an excretion-reliance trade-off might shape the coexistence of competing niche-separated bacterial auxotrophs. We consider four fundamental “phenotypes” associated with excretion and reliance to build an intuition for its consequences for coexistence. Breakdown of obligate mutualism can occur in two ways: either by reversion to independence, where one strain no longer requires the cross-fed resource (low reliance), or by selection for cheaters that benefit from the production of a costly good without paying the cost of its production (low excretion). In our model, we therefore expect that a strain can have a high excretion fraction and a high reliance fraction, serving as a “reliant altruist” that requires high levels of production from its mutualistic partner while simultaneously providing significant benefits (**Fig. 1B, 3D**). Alternatively, a strain with a low excretion fraction and high reliance fraction might grade into a “reliant cheater” phenotype, providing minimal benefits to its partner but depending heavily on cross-fed resources. Given some combination of low reliance and high or low excretion, species can function as either “low-reliance altruists” or “low-reliance cheaters”. While partnering with a reliant altruist might be beneficial for yeast, a bacterial auxotroph with such a trade-off will likely be easily outcompeted by a strain that “cheats” by having lower excretion or moves toward greater independence by reducing reliance, which might ultimately lead to instability or breakdown of the obligate mutualism.

To understand how reliance and excretion influence auxotroph coexistence, we varied the relative reliance and excretion parameters for the two auxotrophs, using T as the reference (**Fig. 3D, S5**). We find that the reliance-excretion parameter relationships that enable coexistence are more complex than what we observed when varying the relative growth rates and amino acid Monod constants, regardless of whether amino acids are cross-fed or externally supplied. During cross-feeding, one bacterial strain can exclude the other only when it is a low-reliance cheater, while coexistence is promoted as long as one auxotroph is either a low-reliance altruist or a reliant cheater relative to its competitor. This observation is consistent with the demands of cross-feeding: along this trade-off, one strain sacrifices more biomass to produce galactose while simultaneously requiring less amino acid (i.e. it is primarily carbon-limited), while the other invests less in galactose production but requires more amino acid for growth (i.e. it is primarily amino acid-limited). Moreover, this trade-off might be expected experimentally: the low-reliance altruist should excrete more galactose, a product of lactose cleavage, because more of its biomass is derived from lactose than from amino acid. Additionally, changing reliance and excretion parameters, unlike growth rate and Monod constants, has dramatic effects on biomass: total system biomass and interguild coexistence frequency varies across the *u*-*l* parameter space (**Fig. S6**). The excretion-reliance trade-off also enables coexistence when amino acids are externally supplied, but over a smaller region of the parameter space (**Fig. S5**). By increasing the effective niche separation of the competing auxotrophs, the reliance-excretion trade-off can therefore support bacterial coexistence across both conditions.

### Intraguild competition for cross-fed resources between bacterial auxotrophs constrains coexistence

While there is significant existing work supporting the idea that coexistence between intraguild competitors can be promoted by a mutualistic partner assuming a high degree of niche separation [Johnson and Bronstein 2019, Hammarlund et al. 2021], it is less clear under what conditions coexistence should be possible when there is incomplete niche separation between our bacterial auxotrophs. Here, we define “incomplete” niche separation as a resource preference hierarchy that means that one auxotroph can use the other’s resource at some reduced rate, while the other can only use a single amino acid. Based on the ability of glycine auxotrophs to convert externally-supplied threonine into intracellular glycine, thus circumventing its glycine auxotrophy [Bertels et al. 2012], this incomplete separation is distinct from models where two strains compete for perfectly substitutable resources. We thus amended our model a third time by replacing the methionine auxotroph (M) with a glycine auxotroph (G) and adding a conversion ratio, *α*, tuning how many units of glycine that one unit of threonine is equivalent to for the glycine auxotroph (**Fig. 4A-B, Methods S2**). When *α* = 1, the two amino acid resources are perfectly substitutable for the glycine auxotroph; for all simulations, we assume a conversion ratio of 0.5 (**Table 1**). We expected that this incomplete niche separation would confer significant disadvantage to the threonine auxotroph (T) under both cross-feeding and conditions of externally supplied resources, but that coexistence might still be possible under a more limited range of parameters.

**Figure 4.**
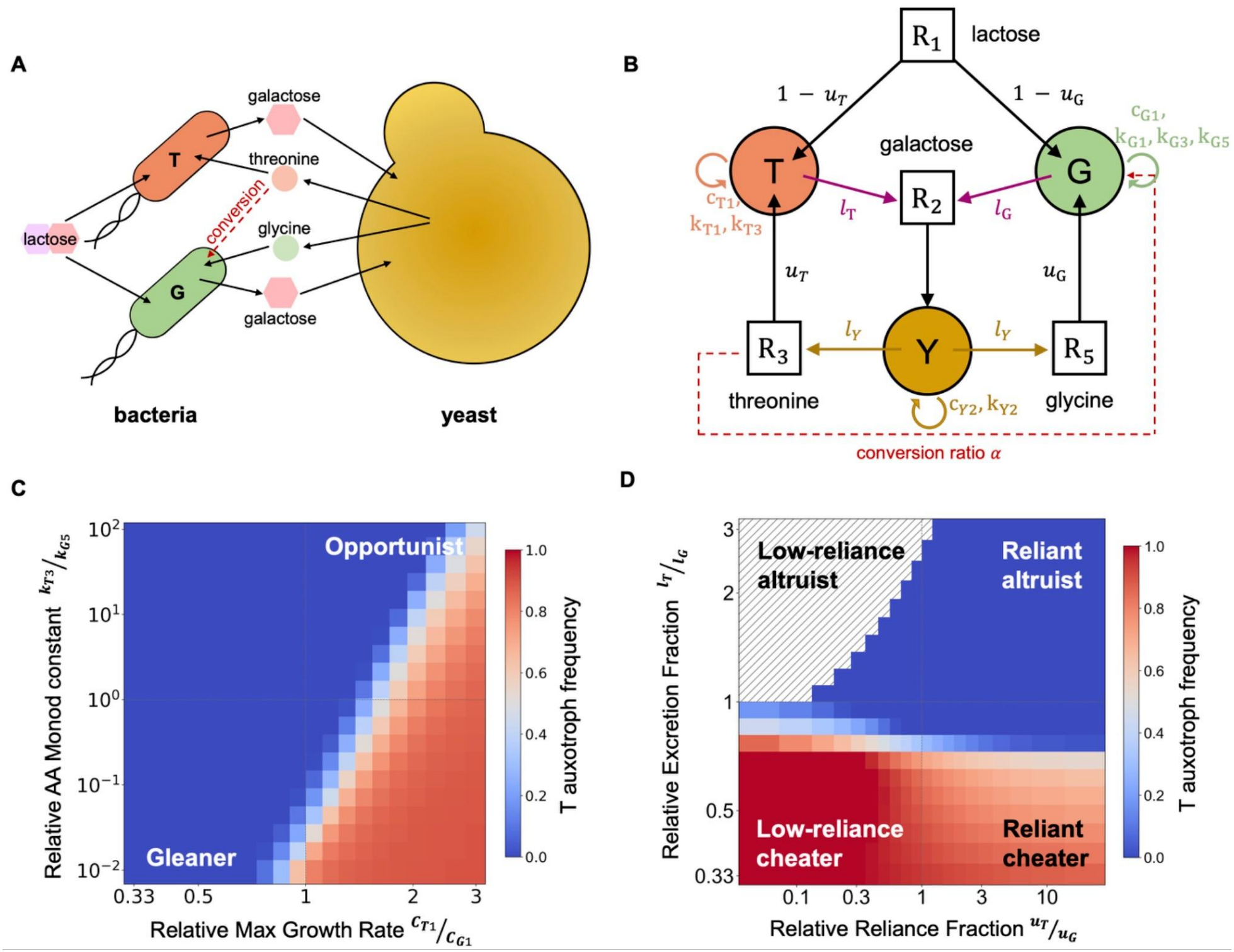
Competition for cross-fed resources shifts the region of the reliance-excretion trade-off that supports coexistence. **A-B**: Schematic representation of the 3 strain system in Eqs. S20-S26. A glycine auxotroph can convert threonine to glycine, resulting in incomplete niche separation. **C:** Heat map of the fraction of the bacterial population composed of the threonine auxotroph (T) during co-culture growth as maximum uptake rates and amino acid half-saturation constants vary. Relative maximum uptake values greater than 1 indicate that T grows faster; relative amino acid half-saturation constants greater than 1 indicate that T requires a higher concentration of its amino acid to reach its maximum growth rate. **D:** Heat map of the fraction of the bacterial population composed of the threonine auxotroph (T) during co-culture growth as excretion and reliance vary. Relative reliance values greater than 1 indicate that T requires more units of amino acid per unit of lactose to generate a single cell; relative excretion values greater than 1 indicate that T excretes a greater percentage of lactose as galactose. The hatched area indicates a region of parameter space where co-culture growth was not possible.

To test this expectation, we varied relative growth rates and amino acid Monod constants in our amended model (**Fig. 4C, S2**). During cross-feeding, we observe a relationship similar to that of the three-species model with niche separation (**Fig. 3B**), with 50/50 coexistence possible along the gleaner-opportunist trade-off boundary. Unsurprisingly, however, competitive exclusion of the threonine auxotroph is more likely. However, as in the TMY model (**Fig. 3, S3, Methods S1**), these parameters still have no effect on total biomass or bacteria-yeast coexistence: regardless of whether G excludes T or they coexist, total bacterial biomass and yeast biomass remain constant (**Fig. S3**). When amino acids are externally supplied, there are regions of parameter space where T can exclude G successfully, but it requires a large growth rate advantage in order to do so, and the total space of coexistence is reduced relative to the cross-feeding model (**Fig. S2**).

Incomplete niche separation also has significant consequences for the reliance-excretion trade-off (**Fig. 4D, S5**). In the regions of parameter space where co-culture growth with yeast is possible, the threonine auxotroph, the bacterial strain with the more constrained niche, can only coexist or exclude the glycine auxotroph if it has a higher reliance fraction (**Fig. 4D**). Collapse of the mutualism is inevitable when T has higher excretion and lower reliance than G (**Fig. 4D**). In this case, high amino acid reliance by G reduces the pool of threonine available to T without sufficient increases in available galactose due to the low excretion of G, thus limiting co-culture growth with yeast. As in the case of the niche-separated model, biomass production depends on excretion and reliance parameters, rather than on growth rates and Monod constants. Biomass peaks near the collapse boundary (**Fig. S6**), recalling the two-species TY model (**Fig. 2**). When resources are externally supplied (**Fig. S5**), coexistence is enabled by the same trade-off shape between excretion and reliance parameters, but over a smaller region of parameter space.

Together, our results indicate that the trade-off between reliance and excretion determines not only for the yield and stability of a two-species mutualism (**Fig. 2**), but also the intraguild coexistence of bacterial auxotrophs with varying niche overlap in our three-species system (**Fig. 3-4**). In the pairwise system, mutualistic reliance sets the minimum excretion needed to sustain cross-feeding, constraining the stability of our yeast-bacteria mutualism. Moreover, in both of our three-species models, excretion and reliance jointly govern intra- and interguild coexistence. Excretion and reliance therefore shape both the qualitative outcomes of mutualism (growth or collapse) and its quantitative outcomes (biodiversity and biomass). We next asked how these two quantitative outcomes relate to one another.

### Reliance-excretion trade-off modulates biodiversity and total biomass

In our pairwise system, bacteria dominate at maximal biomass, with total biomass increasing near the collapse boundary largely as a result of rising bacterial frequency. However, it is not obvious how yeast biomass might change in our three-species system as a function of intraguild coexistence or exclusion. Predicting both strain-specific and community-level biomass is especially important for connecting our model to experimental work, where total mutualistic yield is often used to measure the strength of a mutualism and to assess evolutionary changes in interaction strength [Venkataram et al. 2023]. In multi-mutualist systems, however, total yield may depend not only on the strength of interactions between guilds but also on coexistence within them, making interaction strength difficult to infer from yield alone. We therefore sought to understand how intraguild coexistence shaped community biomass. To address this, we compared the equilibrium biomass of bacteria and yeast in both the TMY (niche separated) and TGY (incompletely separated) models based on simulations where only reliance and excretion parameters were varied between the bacterial strains and all system other parameters were held constant (**Fig. 5**). Yeast biomass was highest when the two bacterial strains were at similar relative abundances, and trended toward its minimum when one auxotroph was competitively excluded (**Fig. 5**). This pattern likely arises because intraguild coexistence of auxotrophs requires at least one strain to excrete galactose at a high rate (i.e. to act as a “low-reliance altruist”), which in turn supports more yeast growth. Reliance and excretion therefore link biodiversity to biomass: the mutualistic trait combinations that allow bacterial coexistence also favor yeast, so total biomass declines as biodiversity increases, both among bacteria and between yeast and bacteria. Lower total yield in multi-mutualist communities may therefore reflect greater intraguild coexistence rather than weaker interguild mutualism.

**Figure 5.**
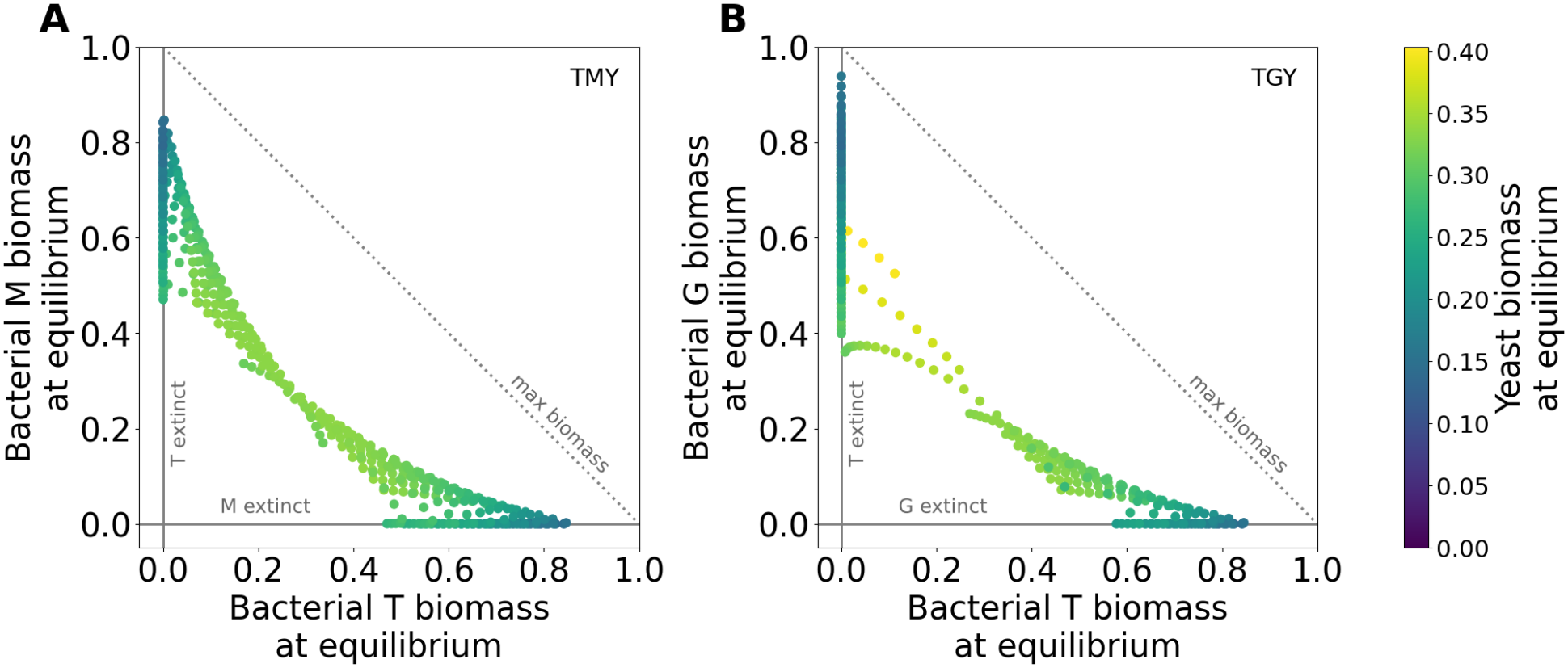
Intraguild coexistence of bacterial strains benefits their cross-feeding partner. **A-B**: Biomass of bacteria T (x-axis) plotted against the biomass of its partner bacteria (y-axis) at equilibrium, coloured by yeast biomass, for (**A**) TMY and (**B**) TGY. Each point is one community from the reliance–leakage map of the same model. For (**A**) TMY, from **Fig. 3C**: reliance *u* = 0. 2 ± *x*, with leakage moved in opposite directions, *l* = 0. 6 ± *y*, for T and M. For (**B**) TGY, from **Fig. 4C**: reliance *u* = 0. 2 ± *x*, with leakage moved in opposite directions, *l* = 0. 6 ± *y*, for T and G.

To further investigate the relationship between biomass and biodiversity, we ran simulations across an additional range of reliance and excretion parameters for both three-species models, again holding all other parameters constant (**Fig. S7**). We found that there is a consistent positive correlation between bacterial coexistence (quantified as Shannon diversity) and yeast biomass. As in the two-species results where bacterial frequency increases with total biomass (**Fig. S1**), and as is consistent with the construction of our model, we observe that low bacterial excretion (i.e. increasing bacterial biomass by minimizing the percentage of lactose converted to galactose instead of biomass) constrains yeast frequency and biomass (**Fig. S8**). Also, in parallel with the two-species result that total biomass is maximized in the region of the excretion-reliance parameter space adjacent to the region of collapse (**Fig. 2B**), we observe maximal biomass adjacent to collapse in the three-species parameter spaces as well (**Fig. S6**). Together, these results show that while coexistence of competing mutualists within a guild may reduce total co-culture biomass, it can also support biodiversity across guilds, which may promote community stability.

## DISCUSSION

Metabolic cross-feeding is common in natural microbial communities [Kost et al. 2023], but it is unclear how these interactions contribute to the biodiversity and stability of complex communities [Palmer & Foster 2022]. This is in part because the strength and outcome of microbial mutualistic interactions, in which two or more species exchange beneficial resources, are dependent on genetic and environmental context. For example, even between obligately cross-feeding strains, competition for a subset of externally supplied resources can shape community outcomes [Jones et al. 2012, Hammarlund et al. 2019, Baichman-Kass et al. 2026]. Mutualisms are also vulnerable to lower- or non-producing cheaters that can weaken or even collapse interactions [Porter et al. 2014, Gore et al. 2009]. Here, we have shown that community collapse in obligate mutualism can be caused not only by low production, but also by excessive reliance on cross-fed resources. The fact that biomass is maximized near this critical excretion-reliance boundary suggests that robust pairwise mutualisms may be vulnerable to collapse when parameters change slightly. By expanding our model to include multi-member mutualist guilds, we see that coexistence within guilds relies on a trade-off between excretion and reliance parameters regardless of the details of intraguild niche separation. Furthermore, increased biodiversity within a guild correlates with increased biomass of the partner species cross-feeding to this guild but lower overall community biomass, providing one example for how biodiversity can be stabilizing for mutualisms [Bronstein 2021].

Our results are consistent with previous work demonstrating that intraguild coexistence can increase during engagement with a cross-feeding partner relative to resource competition for externally supplied resources. [Hammarlund et al. 2021]. We additionally recapitulate the relevance of the classic gleaner-opportunist trade-off [Grover 1990], which usually describes competitive coexistence on a single external resource, in the context of intraguild coexistence on two cross-fed resources. However, we find that the gleaner-opportunist trade-off has no consequences for strain-specific or community-level biomass, while a trade-off between excretion and reliance can both promote intraguild coexistence and shift quantitative outcomes of the system like yield and relative abundance. By employing a reliance fraction to describe the proportion of biomass that bacteria obtain from a resource excreted by yeast (amino acid) compared to an externally supplied resource (lactose), our work allows us to recapitulate any mutualism where partners require non-substitutable mixtures of external and cross-fed resources, such as bacterial-yeast interactions in fermented food communities [Ponomarova et al. 2017, Blasche et al. 2021] or legume-rhizobial bacteria relationships. As a result, we believe that our findings suggest that existing modeling paradigms of cross-feeding, which have primarily focused on the causes and consequences of resource excretion, should be expanded to explicitly include resource reliance. Furthermore, reliance will be relevant to any system where an organism requires multiple essential resources and thus may be a worthwhile addition even to models that omit cross-feeding.

Our work has a number of interesting implications for predicting the dynamics of diverse microbial communities. Firstly, our results demonstrate that reliance may tune the benefits and costs of excretion. For example, in our three-species systems, a low-reliance altruist that excretes more resources can coexist with a reliant cheater that excretes less but relies more on resources excreted by others. This trade-off may provide novel insight into how the Black Queen Hypothesis (BQH) [Morris et al. 2012] influences microbial community assembly and evolution: species that produce resources that other community members require may necessarily be less reliant on deriving their own biomass from mutualistic exchange. Similarly, our findings suggest that there will be hierarchies in the “race to the bottom” when it comes to selection for reduced production as predicted by the BQH. Because auxotrophies differ in how many distinct metabolites can satisfy them, Black Queen dynamics should not act on all gene losses equally. Instead, a strain whose requirement is met by a single compound could be excluded by one whose requirement is met by multiple compounds, as in our TGY model, such that some auxotrophies will be more commonly observed in systems where Black Queen dynamics dominate. Our work also underscores the difficulty of assessing the strength of mutualistic interactions between microbes. While biomass in co-culture is commonly used as a proxy for mutualistic strength in cross-feeding experiments [Wu et al. 2019, Kehe et al. 2021], biomass in our pairwise model is minimized when the mutualistic traits excretion and reliance are greatest. Likewise, when biodiversity in our three-species systems is maximized and thus these interactions should be more stable due to functional redundancy, total biomass is generally low. Total mutualistic yield therefore may be an ineffective metric for assessing strain-level mutualistic traits. However, our observation that biomass is maximized when the both cross-feed amino acid and external resource are exhausted simultaneously suggests that assaying the external metabolites present during different phases of the growth cycle [Ross et al. 2025, Paczia et al. 2012, Ponomorova et al. 2017] could help us understand when a pairwise interaction has approached its trait-based optimum.

Importantly, our study is limited by the construction of our model, in which members of a mutualist guild (bacteria) compete for an external resource (lactose) and excrete a single resource (galactose) to a shared partner (yeast), while the shared partner excretes multiple resources (amino acids) in return. We have assigned bacterial strains a single maximum growth rate, with instantaneous growth determined by the abundance of the external and cross-fed resources, both of which are essential and thus appear as multiplied Monod terms (**Methods**). For this reason, we find that coexistence among bacterial strains within a mutualistic guild depends, in part, on reliance on different cross-fed amino acids from yeast. However, coexistence would be possible on lactose and a single cross-fed amino acid if growth were determined by the minimum of the two resource concentrations rather than their product, with each conferring different maximum growth rates [Johnson and Bronstein 2019]. Additionally, while we find that intraguild coexistence of bacterial auxotrophs supports higher biomass of the shared partner yeast, this result could change if the bacterial strains excreted unique resources [Johnson and Bronstein 2019]. Furthermore, our study does not explore differences among community members in resource-to-biomass yield or production ratios of amino acids excreted by yeast, despite the fact that there will almost certainly be variation in these traits across natural yeast strains [Ponomorova et al. 2017, Correia-Melo et al. 2023]. Further exploration of different cross-feeding models and parameters, including the question of how evolution can modulate cross-feeding [Pauli et al. 2022; Melero-Jiménez et al. 2025], is needed to understand the range of possible community outcomes that result from metabolic exchange.

In summary, our work shows that the extent of both excretion of and reliance on cross-fed resources can be critical for stability across a range of mutualistic scenarios, including the persistence or collapse of a pairwise obligate mutualism and the possibility of intraguild coexistence between competing auxotrophs of varying niche separation. Moreover, through the addition of these parameters, our results contribute a theoretical framework for understanding the roles of biodiversity and biomass for stability in mutualistic systems. We expect that our findings can be integrated into existing models and experimental systems to predict the dynamics of real cross-feeding communities ranging from human-managed fermentations to plant-microbe associations.

## METHODS

### Model description

We constructed a variant of a consumer-resource model quantifying the concentrations of two interacting microbial species, bacteria and yeast, and three resources. The biomass of bacteria increases via multiplied Monod growth on its provided carbon source (lactose) and its required amino acid, while biomass of yeast increases through Monod growth on the carbon resource galactose. Bacteria are thus defined by two Monod constants, one for lactose and one for its amino acid, while yeast is assigned a single Monod constant for its growth on galactose. In addition to relevant Monod constants, our model is constructed such that each microbial strain *i* has intrinsic maximal uptake rate *c*_*i*_, yield *Y*_*i*_, and an excretion/leakage fraction 1 − *l*_*i*_. Excretion modifies the total biomass yield so that the amount of biomass produced per unit of resource decreases with increasing *l* _*i*_, representing the loss of assimilable carbon or nitrogen to waste excretion.

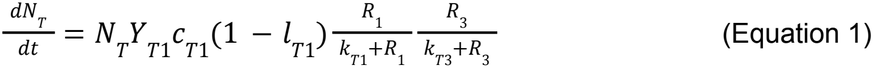

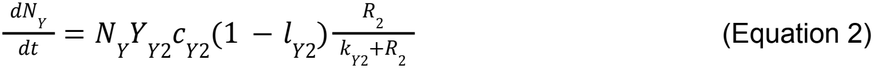

Resources can either be externally supplied or (in the case of galactose or amino acids) generated proportionally to the growth of the relevant microbial strain; depletion is always proportional to microbial growth. Importantly, depletion of lactose and amino acid by bacteria are modified by a reliance fraction *u*_*i*_, where very large values of *u*_*i*_ indicate that the production of a single unit of bacterial biomass requires proportionally more units of amino acid than lactose. The mutualistic dependence of the two strains can thus be manipulated by supplying varying initial concentrations of the three resources and/or changing the values of *u* _*i*_ and *l*_*i*_.

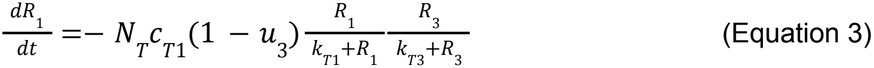

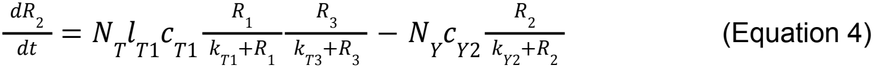

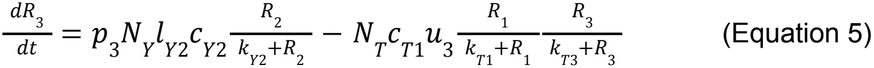

To understand intraguild competition in our cross-feeding mutualism, we add additional bacterial strains and relevant amino acids; in general, these equations are governed by the same fundamental parameters and assumptions as the two-strain, three-resource model (**Methods S1-2**).

### Analytical model evaluation

We begin by evaluating our two-species system of equations analytically. From Eq. 3, we express T’s uptake term, 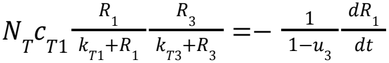, in terms of 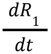. Substituting this into Eq. 4 provides Y’s uptake term, 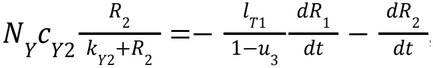, in terms of 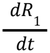 and 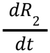. Substituting both uptake terms into Eq. 5 cancels out both biomass terms and uptake parameters (**Discussion S1**). What remains is a linear relationship between 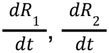 and 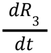:

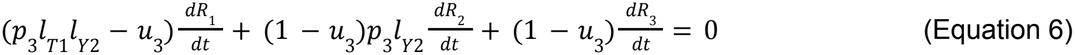

The coefficients are constant, so Eq. 6 integrates to a quantity *K* that is fixed throughout batch growth and is determined by the initial resource concentrations:

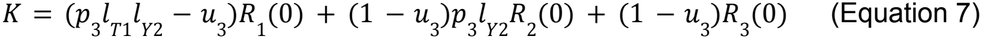

Growth in a single batch necessarily ends when uptake terms for both strains reduce to zero, which requires *R*_2_ = 0 and either *R*_1_ = 0 (complete growth, where all externally supplied lactose is used and threonine remains as a concentration determined by *K*/(1 − *u*_3_)) or or *R*_3_ = 0 (stalled growth/collapse, where lactose remains at a concentration equivalent to *K*/(*p*_3_ *l*_*T*1_ *l*_*Y*2_ − *u* _3_)) (**Discussion S1**).

At the boundary between complete growth and collapse, *K* = 0, allowing us to solve for the threshold *u* _3_:

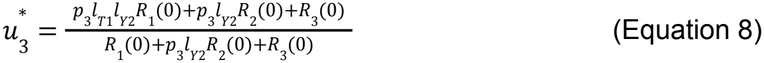

T uses up the lactose (successful growth) when 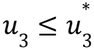 and stalls (collapse of the mutualistic system) when 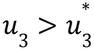.

The biomass equations contain the same uptake terms as those present in the resource equations, so *dN*_*T*_ /*dt* (Eq. 1) and *dN* _*Y*_ /*dt* (Eq. 2) are also constant multiples of the resource rates. Integrating over the batch (**Discussion S2**), with *R*_2_ = 0 at the end, gives the biomass produced:

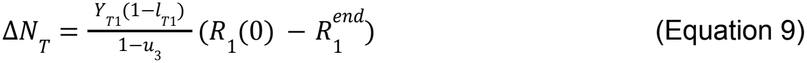

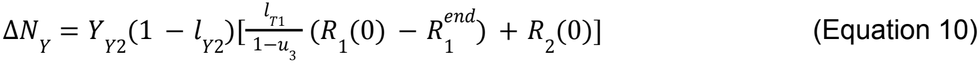

where the amount of lactose consumed is:

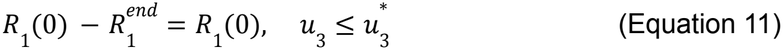

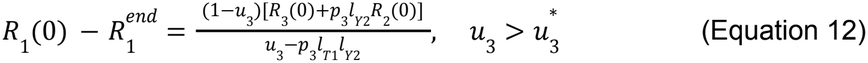

Thus, the summary of biomass increase across all potential growth states should be:

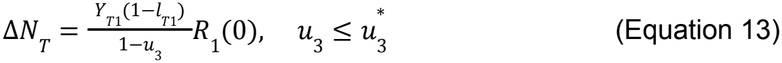

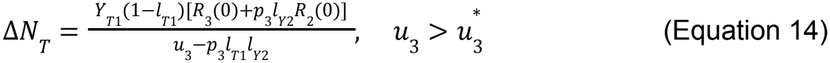

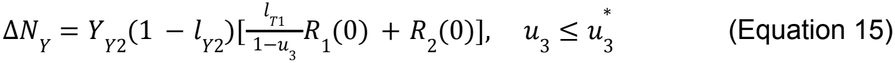

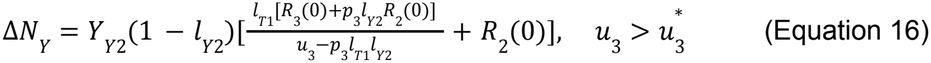

### Model simulations and simulation analysis

To predict the biomass of both yeast and competing bacterial auxotrophs over time, we numerically solved our ordinary differential equations in Python v 3.12 with the solve_ivp function from scipy v. 1.16.3 using the LSODA solver. Serial transfer was performed by carrying 10% of the population to fresh medium with renewed resources (i.e. a population dilution factor of 10) and iterating until populations at the end of each cycle were stable between transfers. We defined this equilibrium transfer as the first point at which biomass and relative abundances changed less than 1e-6 between subsequent transfers. During integration, the relative and absolute tolerances were set to 1e-8 and 1e-11, respectively, and the timestep was set to 48. For all simulations, initial concentrations of all microbial strains were set to 0.01, while lactose was supplied at a concentration of 1, galactose was absent from the initial media, and relevant amino acids were provided at trace levels of 1e-4. Default parameter values and their biological significance are available in **Table 1**.

We evaluated the coexistence of competing bacterial auxotrophs by imposing two different trade-offs: (i) a gleaner-opportunist trade-off, where we simultaneously changed the relative half-saturation constants for each amino acid and the relative maximum uptake rates, and (ii) a reliance-excretion trade-off, where we changed the relative reliance and excretion fractions. Relative values for each parameter were calculated by shifting the parameter values of the threonine auxotroph and its competing partner in opposite directions by adding or subtracting equivalent values (i.e. *c*_*i*_ 0.8 ± x, *u*_*i*_ = 0.2 ± x, *u*_*i*_ = 0.6 ± y; for the half-saturation constant, *k*_*i,j*_ = 0.001 × q for one strain and 0.001 / q for the other). Different starting points were chosen in order to maximize the total parameter space explored. For each simulation, we quantified the relative frequency of the competing auxotrophs using their equilibrium abundance by dividing the density of the threonine auxotroph by the total bacterial biomass. Relative abundance values greater than 0.5 indicated that the threonine auxotroph was more abundant. Simulations were performed for a three-species system where the threonine auxotroph competed either with a methionine auxotroph (niche separation) or a glycine auxotroph (incomplete niche separation).

In some simulations, the co-culture could not meaningfully grow even in the initial cycle (i.e. extinction was not occurring over successive cycles because growth was possible but insufficient to offset dilution rate). Instead, while there was a non-zero steady-state biomass, we observed that strains were not able to metabolize >1% of externally supplied lactose and that total equilibrium biomass remained low (<0.01). We categorized these simulations as collapse and did not calculate relative abundances. The exact value of this collapse threshold was largely arbitrary because, in all cases where we observed that equilibrium biomass rose above 0.01, the total biomass exceeded 0.5.

Finally, to test how the reliance-excretion trade-off shaped strain- and community-level biomass, we used our simulation data from the reliance-excretion trade-off heatmaps to quantify the impact of auxotroph coexistence on biomass. Any simulations where co-culture growth was not possible (i.e. collapse) were omitted. We also tested the relationship between coexistence and biomass across a wider range of parameter space by running simulations where reliance and excretion for bacterial auxotrophs were varied with one of three methods, while keeping the other parameters and initial conditions fixed: (i) varying both strains in opposite directions as previously described, (ii) varying only T strain parameters (*u*_*T*_ =*u*_0_ + *x, l*_*T*_ = *l* _0_+ *y*), or (iii) varying the parameters of the partner bacterial strain (*u*_*partner*_ = *u* _0_+ *x, l*_*partner*_ = *l*_0_ + *y*).

Starting points for each parameter were chosen to keep the highest and lowest values within a reasonable range; for example, for *u*_0_ = 0. 1, 0. 2, 0. 3, the varied value *x* ranged from ± 0. 09, ± 0. 18, or ± 0. 27, respectively. For *l*_0_ = 0. 3, 0. 45, 0. 6, 0. 75, *y* values ranged from ± 0. 20, ± 0. 35, ± 0. 30, or ± 0. 15, respectively, to keep the general excretion range within [0. 1, 0. 9]. To understand the relationship between bacterial diversity and yeast biomass, in particular, we grouped our simulation data by bacterial community leakage values to exclude the effect of bacterial leakage on yeast biomass, allowing us to draw inferences based on bacterial coexistence alone. Bacterial community leakage was defined as the mean of bacterial leakage weighted by biomass, expressed as 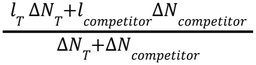. A linear regression was fit to the data corresponding to each leakage group, where Shannon’s diversity *E*_*H*_ was the independent variable and yeast biomass was the dependent variable. The slope and correlation coefficient were subsequently extracted and compared against community leakage (**Fig. S7**).

### Bacterial strains

The Δ*galK ΔmetB* strain has been previously described [Harcombe et al. 2018]. Additional *E. coli* strains are derivatives of a K-12 MG1655 strain obtained from the *E. coli* Genetic Resource Center. The galactose secretion phenotype was generated by knocking out the function of the Δ*galK* gene via lambda red recombineering with the pSIM6 plasmid as previously described [Thomason et al. 2023]. The *ΔgalK* gene in MG1655 was replaced with the kanamycin cassette amplified with appropriate homology from the pKD4 plasmid as described [Datsenko and Wanner 2000]. To resensitize the MG1665 *ΔgalK* strain to kanamycin, we transformed the strain with the pCP20 plasmid, which facilitates excision of the kanamycin cassette via flippase [Cherepanov et al. 1995]. After excision, we generated a library of single amino acid auxotrophs with the pSIM6 plasmid by knocking out the function of genes whose products catalyze the terminal step of the corresponding amino acid synthesis pathway: proline (*ΔproC*), threonine (*ΔthrC*), or glutamine (*ΔglnA*). All strains are listed in **Table S1**.

### Media and growth assays

Strains were grown in monoculture in a modified M9 minimal medium with lactose as the sole carbon source, containing 13.9 mM lactose monohydrate, 47.8 mM anhydrous Na_2_HPO_4_, 22 mM KH_2_PO_4_, 18.7 mM NH_4_Cl, 2.0 mM MgSO_4_, 0.253 mM CaCl_2_, 8.56 mM NaCl, metal mix (257 uM EDTA, 80.9 uM H_3_BO_3_, 78.2 uM ZnSO_4_ · 7H_2_O, 54.0 uM FeSO_4_ · 7H_2_O, 25.3 uM MnCl_2_ · 4H_2_O, 8.27 uM Na_2_MoO_4_ · 2H_2_O, 6.30 uM CoCl_2_ · 6H_2_O, 6.01 uM CuSO_4_· 5H_2_O, 3.01 uM KI, 0.025% v/v silicone antifoam), and vitamins (694 uM myo-Inositol, 40.6 uM nicotinic acid, 24.3 uM pyridoxine HCl, 14.8 uM thiamine HCl, 10.5 uM Ca pantothenate, 7.29 uM *p*-aminobenzoic acid, 1.02 uM biotin). Strains were grown in this medium with between 0 and 5000 uM of appropriate amino acid added to identify concentrations where growth was limited by lactose depletion rather than amino acid consumption. Assays were completed in 96-well plates with 200uL of media per well inoculated with 1uL of a stationary overnight culture diluted to an OD600 of 0.1. We measured the initial OD600 of each well as well as its final OD600 after 48 hours of growth at 30C without shaking. The maximum OD600 was calculated as the final OD600 subtracted by the initial OD600. For all experiments, we performed three biological replicates per amino acid concentration. Overnight cultures were grown in 5mL of LB Miller broth with 50 ug/mL of kanamycin with shaking at 200rpm at 30C.

### Flux balance analysis simulations and data analysis

Flux balance analysis (FBA) and parsimonious flux balance analysis (pFBA) were performed as previously described [Orth et al. 2010]. Simulations were completed in COBRAPy v. 0.32.1 with the GLPK solver using the existing *E. coli* K-12 MG1655 model iJO1366 (2,583 reactions, 1,805 metabolites, 1,367 genes) [Ebrahim et al. 2013]. We used the iJO366 biomass function (BIOMASS_Ec_iJO1366_core_53p95M) as the objective function and defined an M9 medium under aerobic conditions by setting the lower bounds of the following exchange reactions to 1000 millimoles per gram of dry weight per hour (mmol gDW^-1^ hr^-1^): EX_na1_e, EX_k_e, EX_pi_e, EX_cl_e, EX_nh4_e, EX_so4_e, EX_ca2_e, EX_fe2_e, EX_fe3_e, EX_zn2_e, EX_mn2_e, EX_cu2_e, EX_cobalt2_e, EX_mobd_e, Ex_ni2_e, EX_sel_e, EX_slnt_e, EX_tungs_e, EX_h2o_e, EX_h_e, EX_co2_e, EX_o2_e. Lactose was supplied as the sole carbon and limiting reagent at an uptake rate of 5 mmol gDW^-1^ hr^-1^. The *galK* (b0757) gene was deleted with COBRAPy gene.knockout() to match the background genotype of the amino acid auxotrophs used for *in vitro* studies. Next, we deleted the genes *glnA* (b3870), *thrC* (b0004), *proC* (b0386), or *metB* (b3939) to recapitulate strains with single amino acid auxotrophies for glutamine, threonine, proline or methionine, respectively. Because the glutamine synthetase reaction (GLNS) can be governed by multiple genes, we eliminated flux through this reaction entirely in additional to knocking out the *glnA* gene to produce complete glutamine auxotrophy; no additional constraints were required for the other three genes, since their target reactions are governed by solely by their corresponding genes. Auxotrophy was verified for each strain by simulating growth on lactose minimal media with and without amino acid supplementation.

To determine the concentration of amino acid required to switch auxotrophic growth from amino acid limitation to carbon limitation, we first optimized our wild-type model (MG1655 Δ*galK*) on lactose M9 minimal media, giving us a carbon-limited reference growth rate of 0.9772 hr^-1^. For each auxotroph, the exchange reaction for the relevant amino acid was set to 1000 mmol gDW^-1^ hr^-1^ and the flux through the biomass reaction was constrained to the reference growth rate by fixing the upper and lower bounds. We then performed FBA with the objective function set to the amino acid exchange reaction to identify the optimal amino acid flux required to meet cellular biosynthetic demands. Using these simulations, we calculated the number of amino acid and lactose molecules consumed per gram of dry weight *c*_*i*_ by converting fluxes to biomass-normalized consumption such that 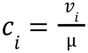, where *v*_*i*_ was the optimal flux through the appropriate exchange reaction in mmol gDW^-1^ hr^-1^ and μ was the cellular growth rate in hr^-1^ (**Table S2**). We then calculated the ratio of the two consumption values by dividing the amount of amino acid consumed per gram of dry weight by the amount of lactose consumed per gram of dry weight, such that values below one indicated greater demand for lactose. This ratio is dimensionless and independent of the number of cells assumed per gram of dry weight.

Finally, we ran pFBA simulations where amino acid uptake was constrained to its identified optimal flux, the objective function was set as the biomass reaction, and total flux through the system was minimized. These simulations provided unique flux distributions from which we could infer the demand carried by each cellular reaction for a given amino acid. We computed demand for an amino acid *m* through reaction *r* in mmol gDW^-1^ hr^-1^ using COBRAPy Metabolite.summary() such that *d* _*r*_ = *v*_*r*_ *S*_*m,r*_, where *S*_*m,r*_ is the stoichiometric coefficient of amino acid *m* in reaction *r* and *v*_*r*_ is the predicted flux through reaction *r* in mmol gDW^-1^ hr^-1^ (**Table S3**). Reactions where *d*_*r*_ was less than −10^−6^ mmol gDW^-1^ hr^-1^ were considered sinks. We summed the demand across all sink reactions for each amino acid and quantified the demand through the biomass reaction as a percentage of total demand.

## DATA AVAILABILITY

All code required to produce the figures is available on GitHub at TBD link. All data generated by experiments and simulations is available in the same GitHub repository.

## ACKNOWLEDGEMENTS

We acknowledge members of the Abreu lab for helpful feedback on the manuscript. We also thank Hyunseok Lee for helpful conversations. Finally, we thank WR Harcombe and the *E. coli* Genetic Resource Center for providing the *E. coli* strains used for experiments.

## SUPPLEMENT

### Supplementary Tables

**Table S1.**
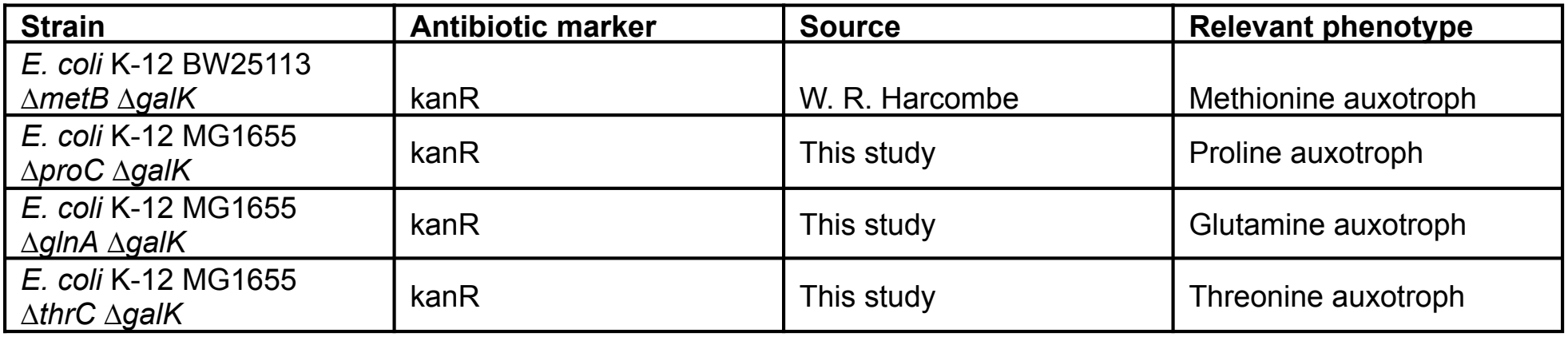
Bacterial strains. Bacterial strains used for experiments. See Methods and Results for additional details.

| **Strain** | **Antibiotic marker** | **Source** | **Relevant phenotype** |
|---|---|---|---|
| *E. coli* K-12 BW25113 Δ*metB* Δ*galK* | kanR | W. R. Harcombe | Methionine auxotroph |
| *E. coli* K-12 MG1655 Δ*proC* Δ*galK* | kanR | This study | Proline auxotroph |
| *E. coli* K-12 MG1655 Δ*glnA* Δ*galK* | kanR | This study | Glutamine auxotroph |
| *E. coli* K-12 MG1655 Δ*thrC* Δ*galK* | kanR | This study | Threonine auxotroph |

**Table S2.** Flux balance analysis indicates that lactose is in greater demand than amino acids regardless of tested auxotrophy. Optimal lactose flux for all auxotrophs was constrained to 5 mmol gDW^-1^ hr^-1^. Amino acid and lactose consumption per gram of dry weight were calculated by dividing the appropriate optimal fluxes by the strain growth rate, which was 0.9772 hr^-1^ for all simulations.

| **Gene knockout** | **Amino acid** | **v<sub>aa</sub> (mmol gDW<sup>-1</sup> hr<sup>-1</sup>)** | **c<sub>aa</sub> (mmol gDW<sup>-1</sup>)** | **v<sub>lactose</sub> (mmol gDW<sup>-1</sup> hr<sup>-1</sup>)** | **c<sub>lactose</sub> (mmol gDW<sup>-1</sup>)** | **Log10 ratio of amino acid consumption to lactose consumption** |
|---|---|---|---|---|---|---|
| glnA | gln__L_c | 1.769 | 1.811 | 5 | 5.117 | -0.451 |
| proC | pro__L_c | 0.2160 | 0.221 | 5 | 5.117 | -1.364 |
| thrC | thr__L_c | 0.5442 | 0.544 | 5 | 5.117 | -0.973 |
| metB | met__L_c | 0.1539 | 0.154 | 5 | 5.117 | -1.521 |

**Table S3.** Amino acids differ in their predicted demand due to their varying roles in biosynthetic pathways. Amino acid sinks were identified using pFBA and quantified as a fraction of total amino acid demand.

| **Gene knockout** | **Amino acid** | **Reaction** | **d<sub>reaction</sub> (mmol gDW<sup>-1</sup> hr<sup>-1</sup>)** | **Reaction function** | **Percentage of total amino acid demand (%)** |
|---|---|---|---|---|---|
| glnA | gln__L_c | PRFGS | 0.43801246 | Fourth step of purine synthesis | 24.7561188 |
| glnA | gln__L_c | GLUPRT | 0.43801246 | First step of purine synthesis | 24.7561188 |
| glnA | gln__L_c | BIOMASS | 0.2571566 | Glutamine incorporated into protein | 14.5342881 |
| glnA | gln__L_c | GMPS2 | 0.23815617 | Synthesis of guanine nucleotides | 13.4603988 |
| glnA | gln__L_c | CTPS2 | 0.15686298 | Synthesis of cytosine nucleotides | 8.86577213 |
| glnA | gln__L_c | IG3PS | 0.09257677 | Histidine biosynthesis | 5.2323658 |
| glnA | gln__L_c | GF6PTA | 0.09233247 | Peptidoglycan and LPS synthesis | 5.21855834 |
| glnA | gln__L_c | ANS | 0.05554626 | First step of tryptophan biosynthesis | 3.13943052 |
| glnA | gln__L_c | ADCS | 0.00065374 | Folate biosynthesis | 0.03694877 |
| proC | pro__L_c | BIOMASS | 0.21601213 | Proline incorporated into protein | 100 |
| thrC | thr__L_c | THRD_L | 0.28390123 | First step of isoleucine biosynthesis | 53.3848692 |
| thrC | thr__L_c | BIOMASS | 0.2478997 | Threonine incorporated into protein | 46.6151308 |
| metB | met__L_c | BIOMASS | 0.15018 | Methionine incorporated into protein | 99.4190861 |
| metB | met__L_c | METAT | 0.00087751 | S-adenosylmethionine (SAM) synthesis | 0.58091394 |

### Supplementary Figures

**Supplementary Figure 1.**
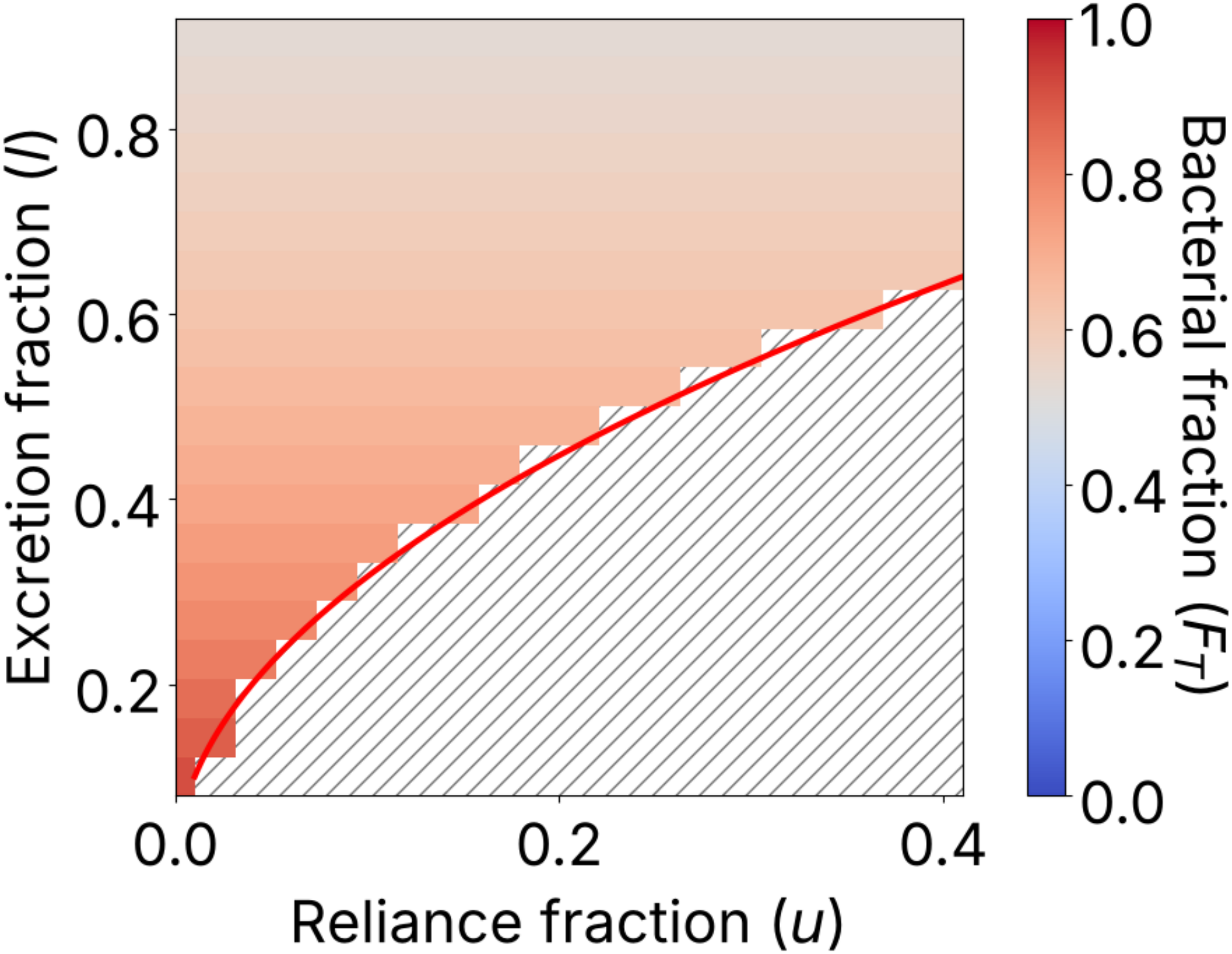
Bacterial frequency changes with only the excretion fraction parameter and reaches maximum at the collapse boundary. This plot is identical to the biomass heat map in **Fig. 2B**; here, the heat map shows bacterial frequency rather than total system biomass.

**Supplementary Figure 2.**
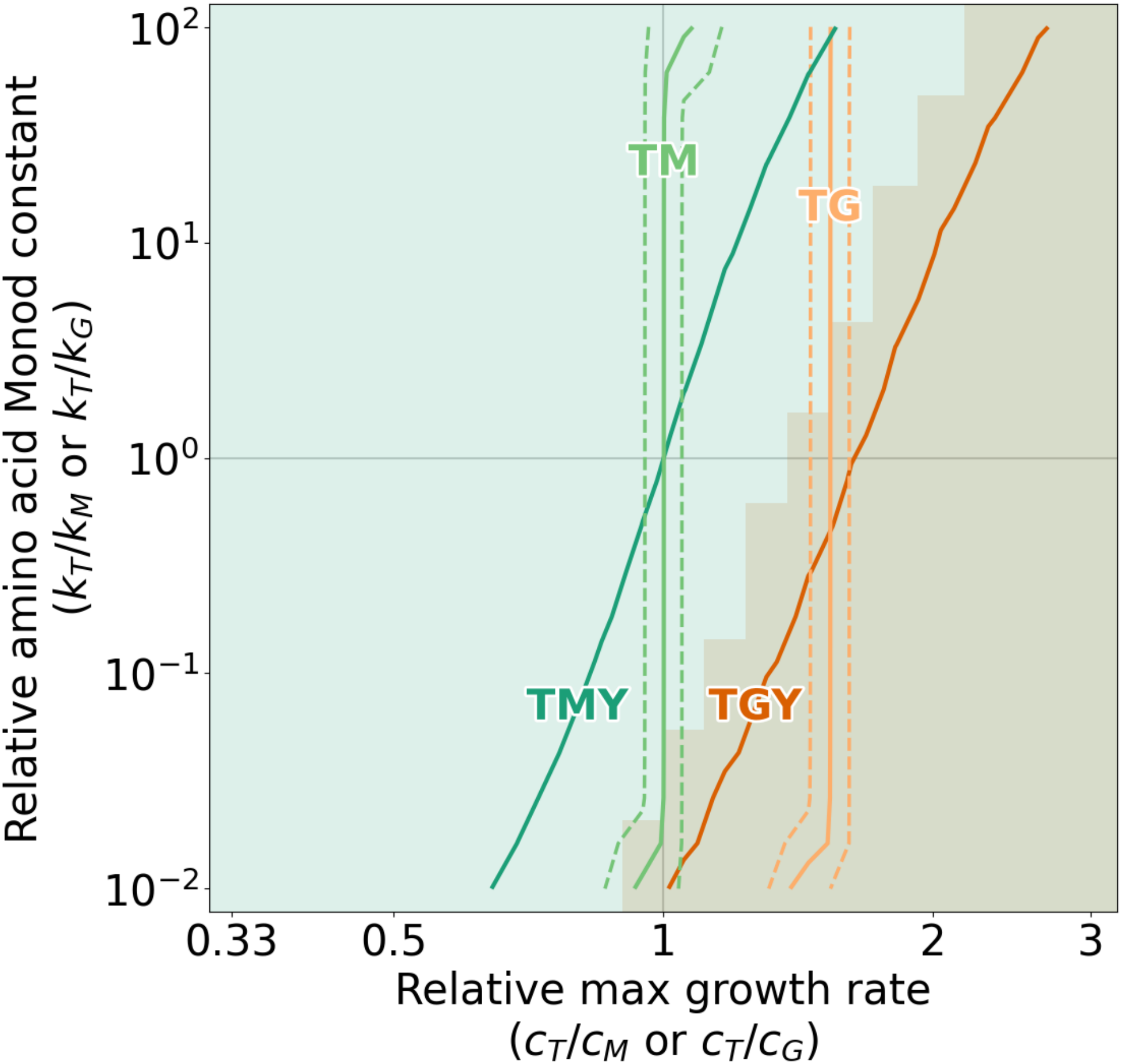
Summary of the gleaner-opportunist trade-off heatmap across TMY, TGY, TM, TG models. Resource supplies are kept consistent, with TMY, TGY model initial concentrations at [1, 0, 1e-4, 1e-4] and TM, TG model initial concentrations at [0.8, 0, 0.1, 0.1]. *f*_*T*_ =0.5 lines are shown for all model types as solid lines.Coexistence areas (<99% one strain) are shaded for TMY and TGY model, and marked with dashed lines for TM and TG model. In the TM and TG models, the gleaner-opportunist trade-off is not as apparent as might be expected given that it is usually associated with resource competition, because while bacteria compete for lactose in the TM and TG models, their ability to coexist is facilitated by private amino acids that are externally supplied.

**Supplementary figure 3.**
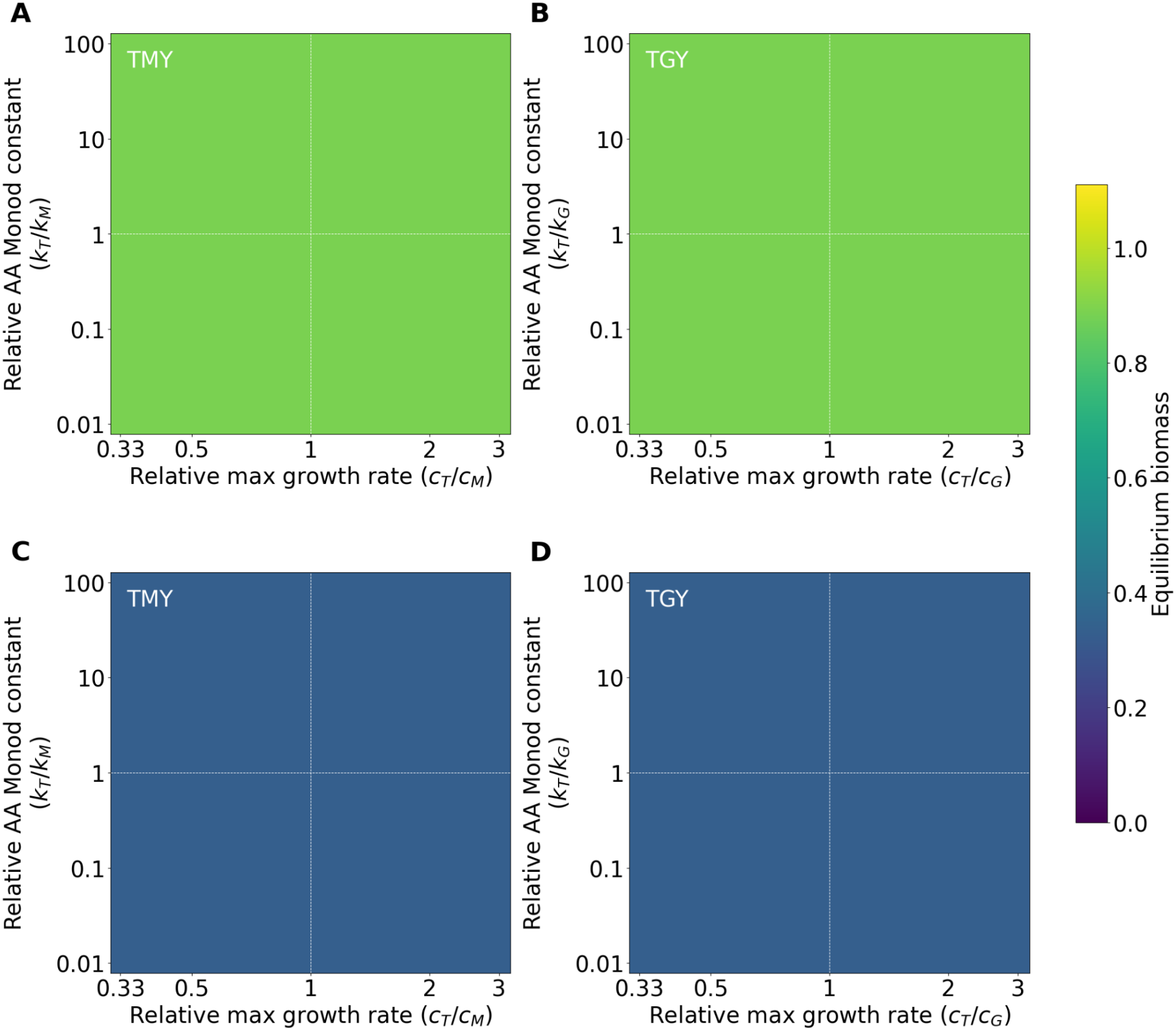
In the three-species TMY and TGY models, Max growth rates and Monod constants have no effect on total biomass or yeast biomass. **A-B**: Equilibrium total system biomass of TMY and TGY models does not change as relative growth rates and amino acid Monod constants are varied. **C-D**: Equilibrium yeast biomass in the same simulations.

**Supplementary figure 4.**
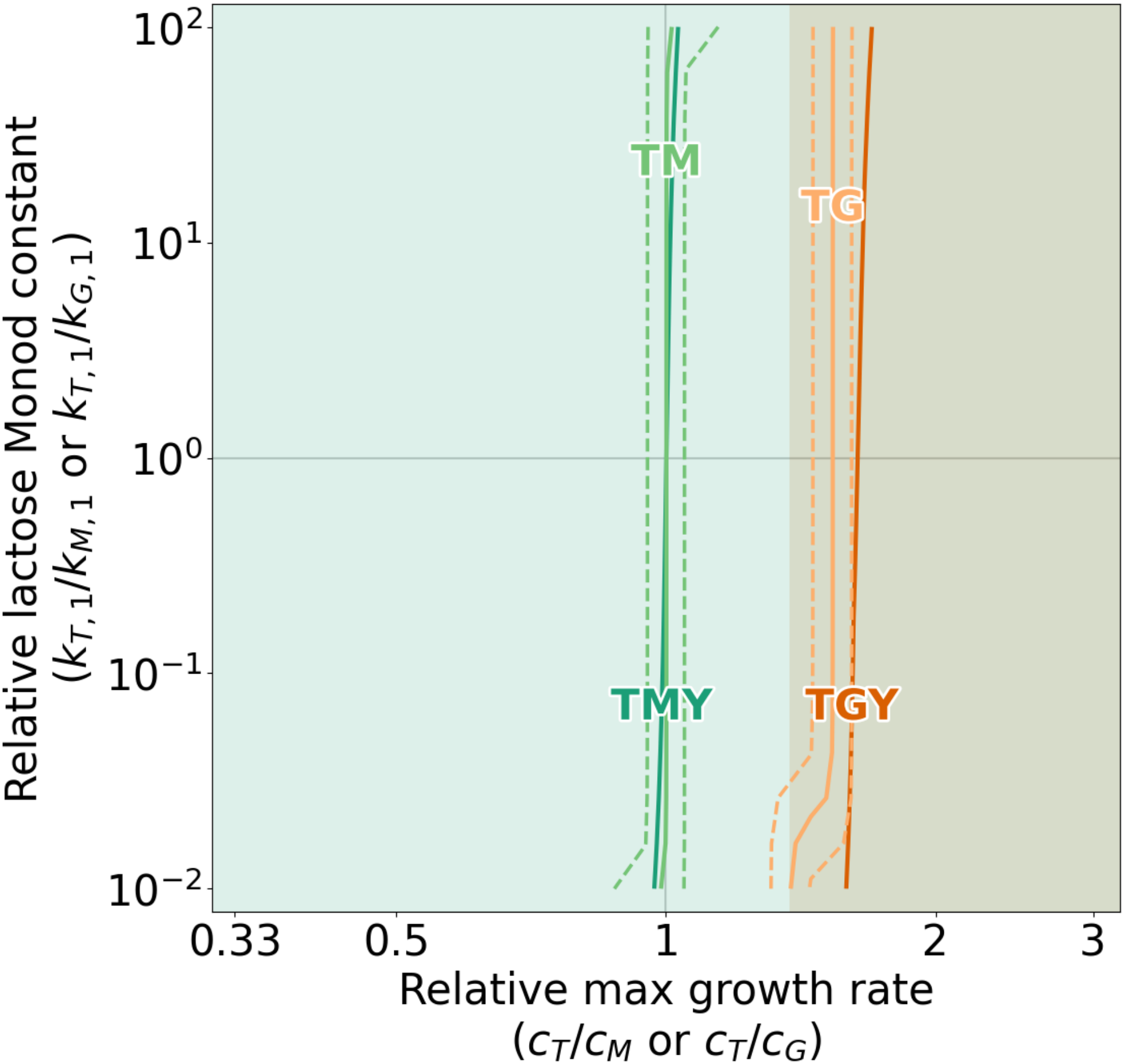
Summary of the gleaner-opportunist trade-off heatmap across TMY, TGY, TM, TG models: lactose Monod constants instead of amino acids. Resource supplies are kept consistent, with TMY, TGY model initial concentrations at [1, 0, 1e-4, 1e-4] and TM, TG model initial concentrations at [0.8, 0, 0.1, 0.1]. *f*_*T*_ =0.5 lines are shown for all model types as solid lines. Coexistence areas (<99% one strain) are shaded for TMY and TGY model, and marked with dashed lines for TM and TG model.

**Supplementary figure 5.**
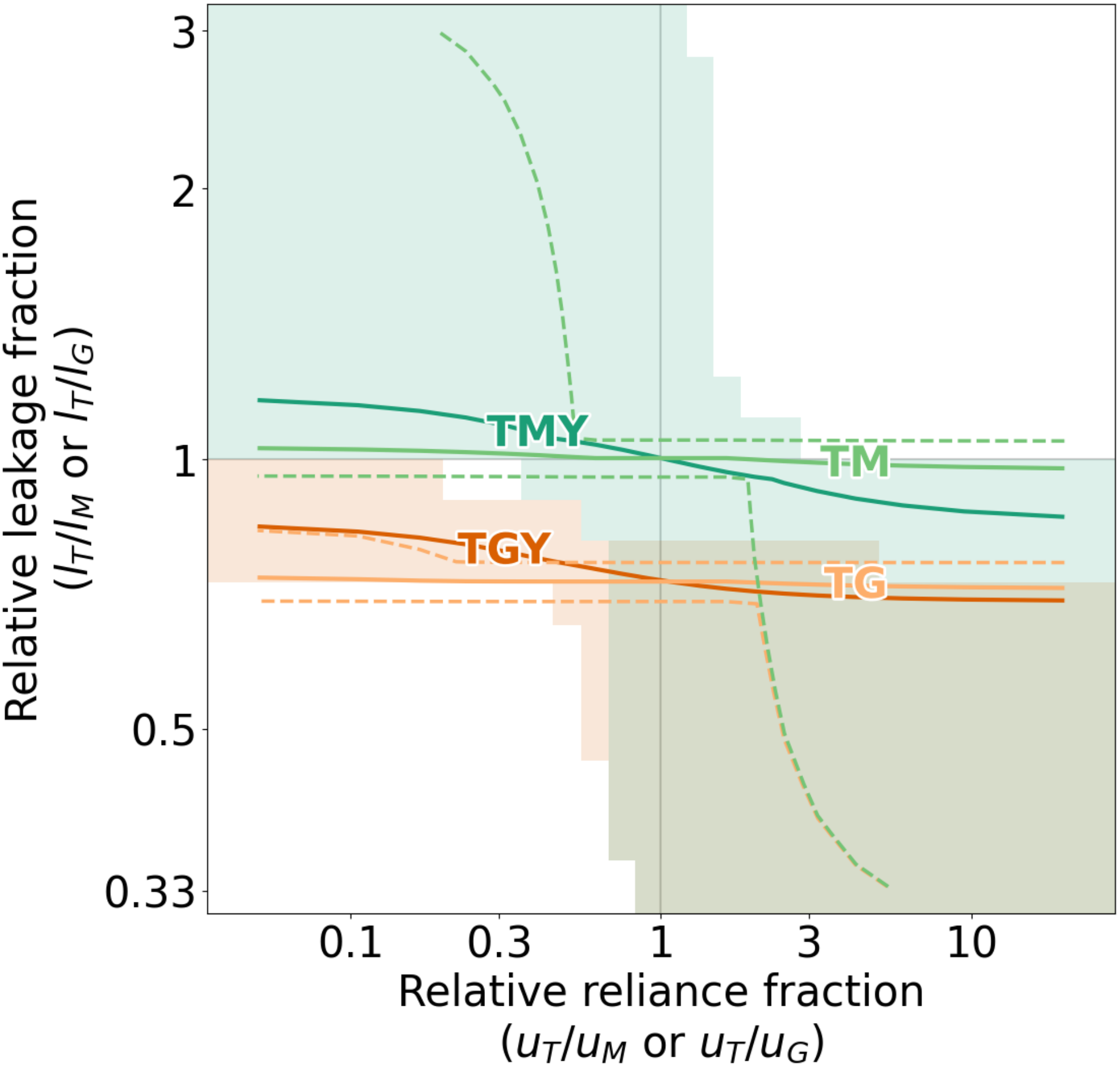
Summary of the reliance-excretion trade-off heatmap across TMY, TGY, TM, TG models. Resource supplies are kept consistent, with TMY, TGY model initial concentrations at [1, 0, 1e-4, 1e-4] and TM, TG model initial concentrations at [0.8, 0, 0.1, 0.1]. *f*_*T*_ =0.5 lines are shown for all model types as solid lines. Coexistence areas (<99% one strain) are shaded for TMY and TGY model, and marked with dashed lines for TM and TG model.

**Supplementary Figure 6.**
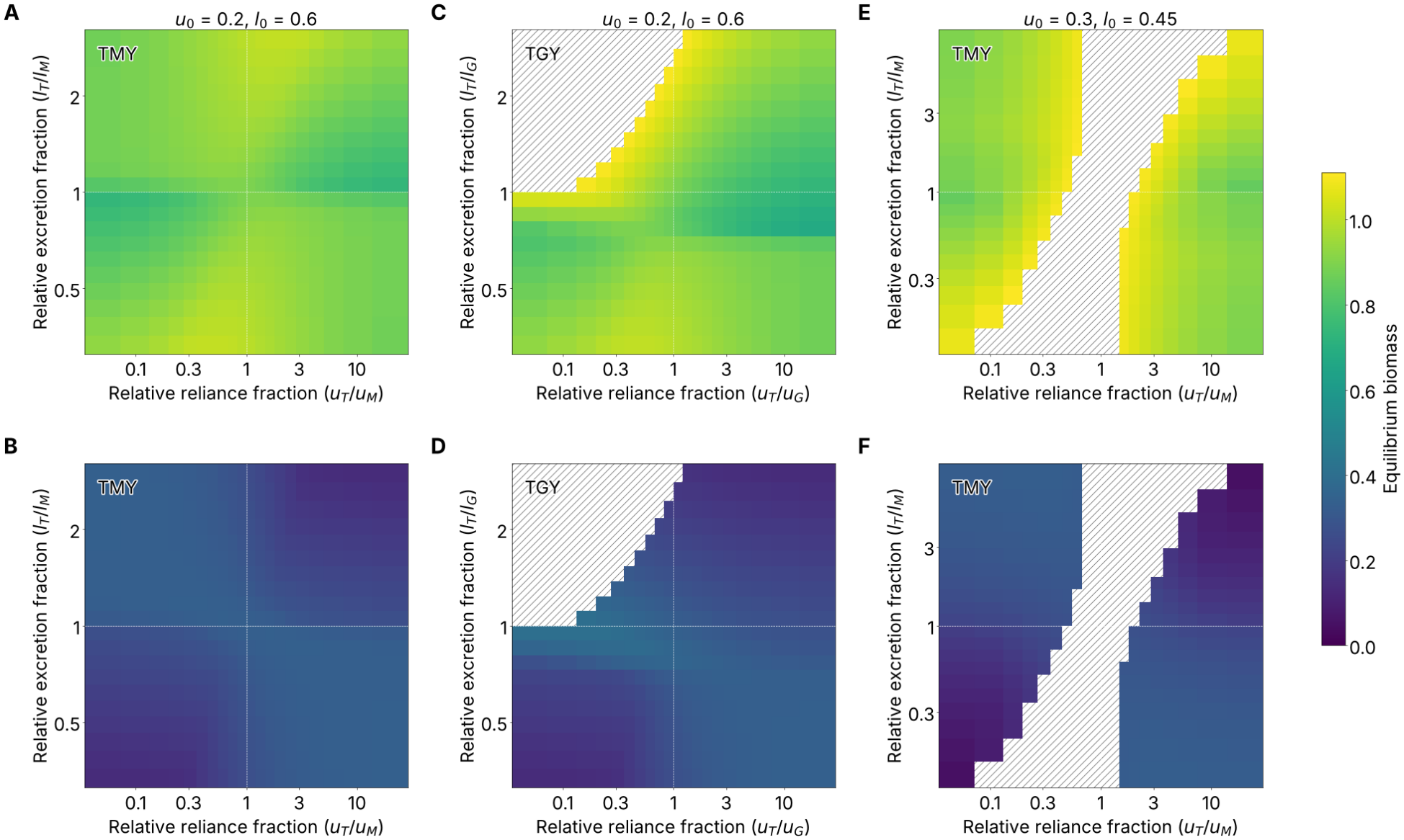
Biomass peaks near system collapse; bacterial coexistence favors yeast. The biomass data from heat maps in main text Figs. 3-4 is shown in panels **A-D**. Simulations that include a collapse region of the TMY model are shown in panels **E-F**. Top row (**A, C, E**): total biomass; bottom row (**B, D, F**): yeast biomass.

**Supplementary Figure 7.**
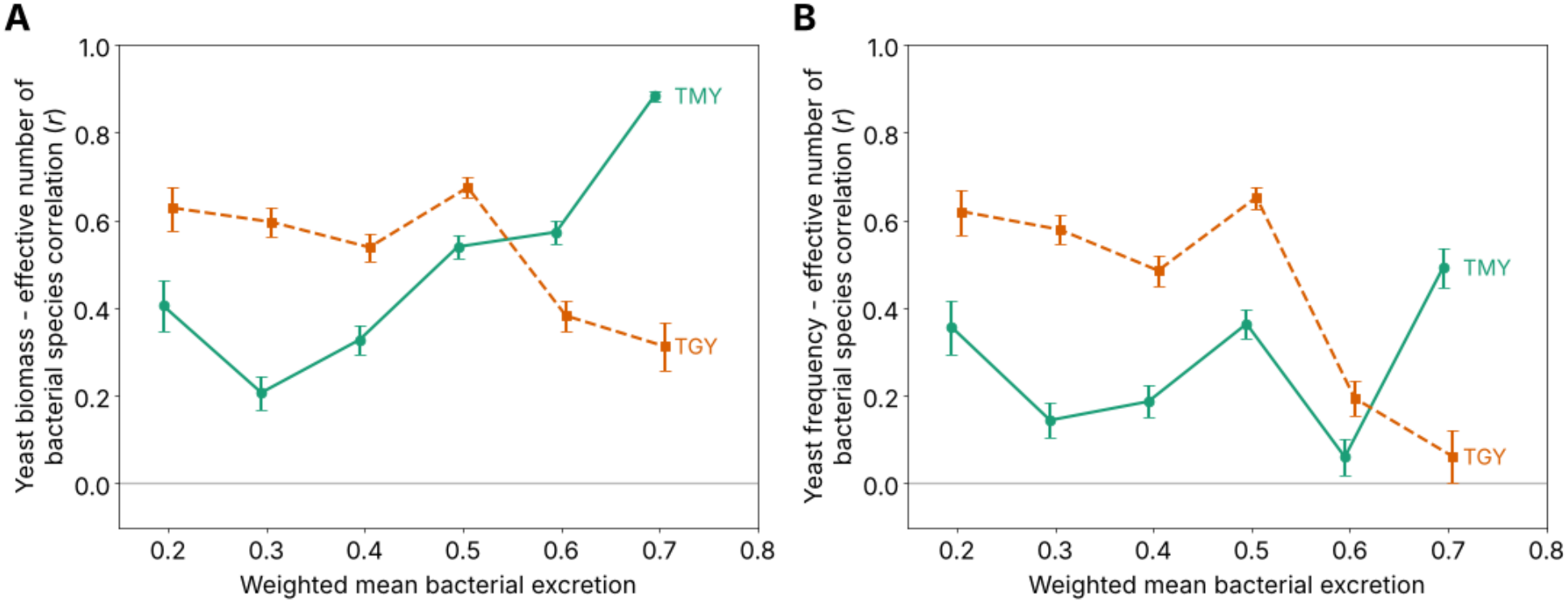
Yeast biomass and frequency are positively correlated with bacterial diversity. Data points of the top figure come from **Figs. S9, S10**, in addition to simulations where u and l span a range of (0.01, 0.57) and (0.1, 0.9). Linear regression lines and slopes are shown, indicating positive correlation between yeast abundance and bacterial evenness. The bottom figure is simulated with the same data with all three variation methods, but grouped by community leakage. The x-axis, weighted mean bacterial excretion, is calculated as 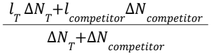 (excretion weighted by biomass). The y-axis shows the linear regression lines for the points under each excretion level, where the x-axis is bacterial Shannon’s H. Positive values on the y-axis suggest correlations between bacteria diversity and yeast biomass and frequency. 95% confidence intervals are shown as the error bar.

**Supplementary Figure 8.**
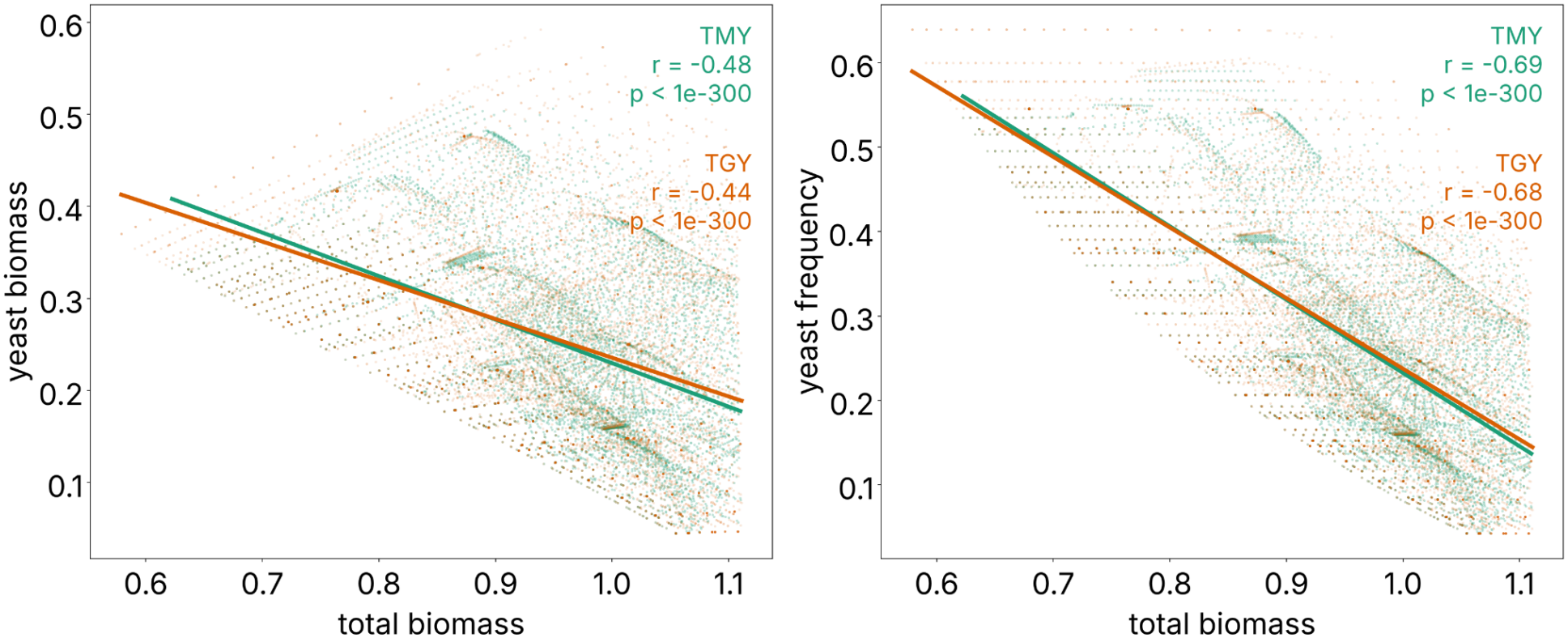
Yeast biomass and frequency are negatively correlated with total biomass. Data points come from **Figs. S9, S10**, in addition to simulations where u and l span a range of (0.01, 0.57) and (0.1, 0.9). Linear regression lines and slopes are shown, indicating negative correlation between yeast growth and total population size.

**Supplementary Figure 9.**
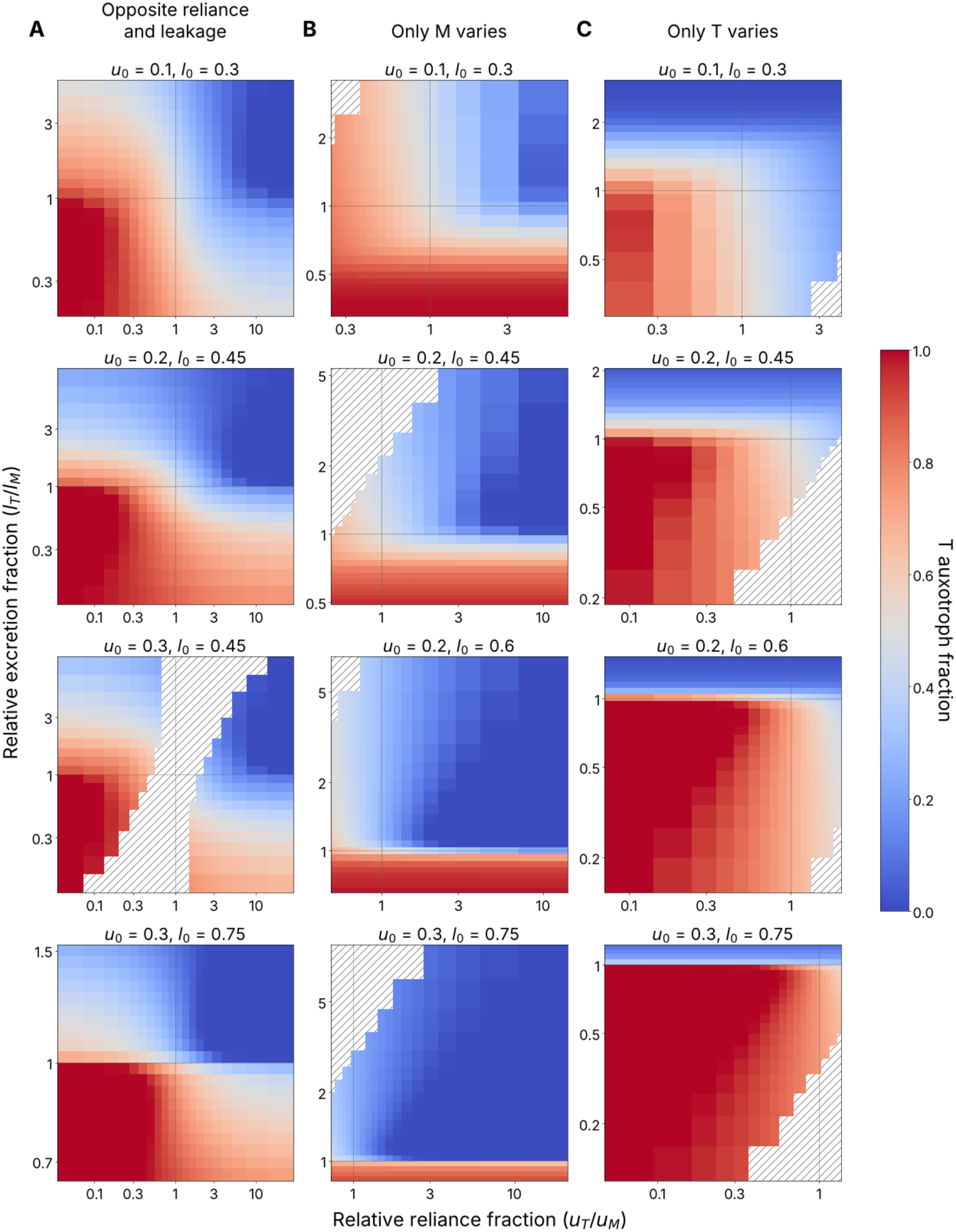
Regardless of method to vary reliance and excretion in the TMY model, trade-off enables coexistence. Method A: both strains’ parameters are varied in opposite directions (*u*_*T*_ =*u* _0_+ *x, u*_*M*_ = *u*_0_ − *x*; *l*_*T*_ = *l*_0_ + *y, l*_*M*_ = *l*_0_ − *y*). Method B: only T strain’s parameters are varied (*u*_*T*_ =*u*_0_ + *x*; *l*_*T*_ = *l*_0_ + *y*). Method C: only M strain’s parameters are varied (*u*_*M*_ =*u*_0_ + *x*; *l*_*M*_ = *l*_0_ + *y*). *x* and *y* can be either positive or negative, so the parameters vary in both directions.

**Supplementary Figure 10.**
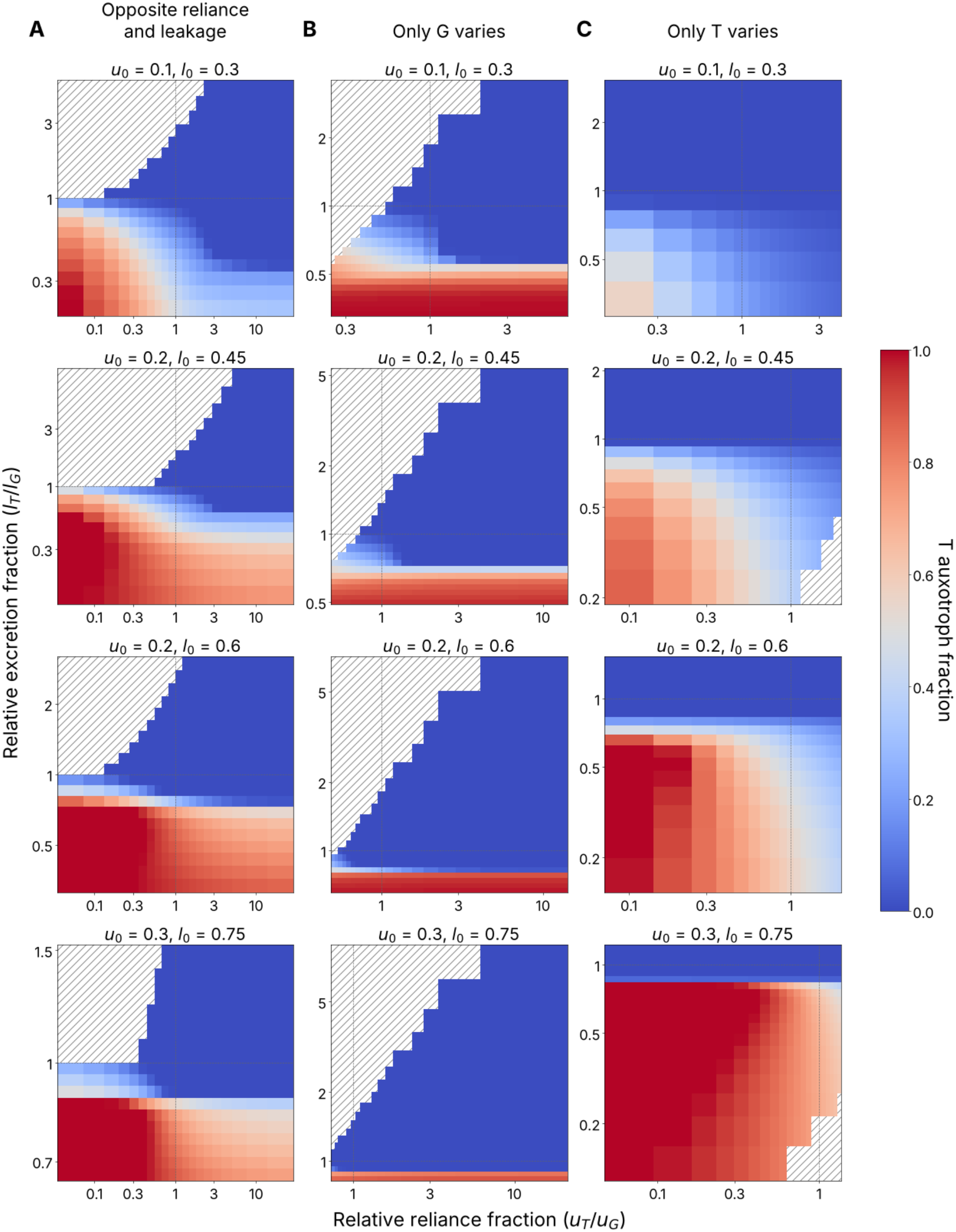
Regardless of method to vary reliance and excretion in the TGY model, trade-off enables coexistence. Method A: both strains’ parameters are varied in opposite directions (*u*_*T*_ =*u*_0_ + *x, u*_*G*_ = *u*_0_ − *x*_0_; *l*_*T*_ = *l* _0_ + *y, l*_*G*_ = *l*_0_ − *y*). Method B: only T strain’s parameters are varied (*u*_*G*_ =*u*_0_ + *x*; *l* _*T*_ = *l*_0_ + *y*). Method C: only G strain’s parameters are varied (*u*_*G*_ =*u*_0_ + *x*; *l*_*G*_ = *l*_0_ + *y*). *x* and *y* can be either positive or negative, so the parameters vary in both directions.

### Supplementary Discussion

#### Supplementary Text 1: Derivation of the analytical expression for

The T–Y batch model is

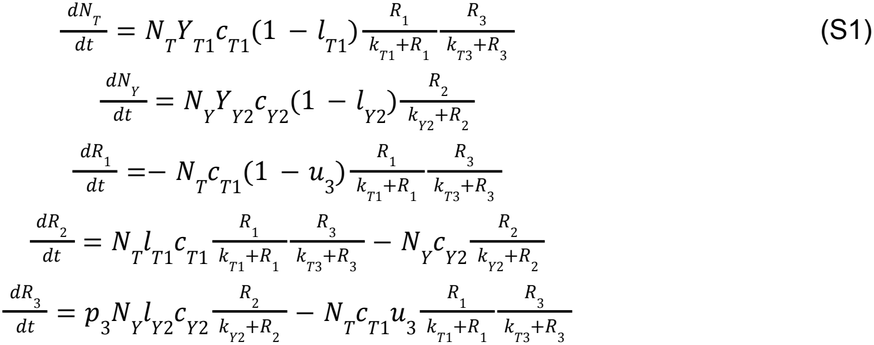

Here *u*_3_ is the reliance of bacteria strain T on the amino acid: the fraction of its uptake drawn from threonine (*R*_3_). The remaining 1 − *u*_3_ is drawn from lactose (*R*_1_). We call 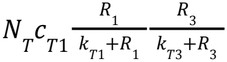 T’s uptake term and 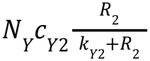 Y’s uptake term.

The *R*_1_ equation contains T’s uptake term, which can be expressed as:

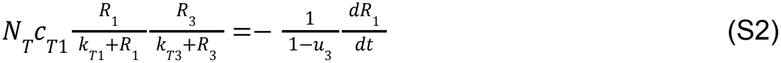

Substituting (S2) into the *R* equation and solving for Y’s uptake term,

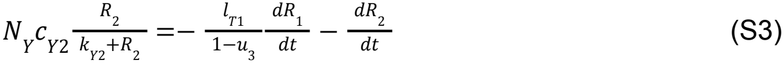

Putting (S2) and (S3) into the *R* equation,

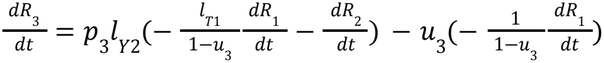

Multiplying through by 1 − *u* and collecting the three derivatives,

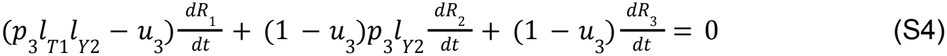

The coefficients in (S4) are constants, so its left side is an exact time derivative. The combination below therefore keeps its initial value for the whole batch:

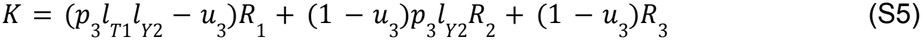

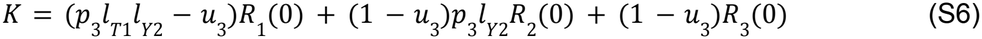

Ending states:

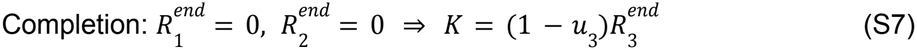

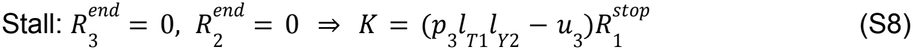

#### Supplementary Text 2: Derivation of analytical expressions for Δ*N*_*T*_ and Δ*N* _*Y*_ (biomass) in the TY model

Factoring the Monod term out from (Equation 1):

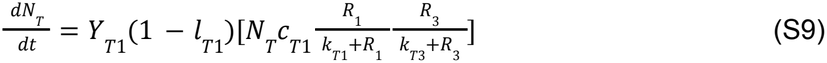

Replacing the bracket with (S2):

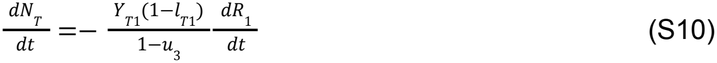

Factoring the Monod term out from (Equation 2):

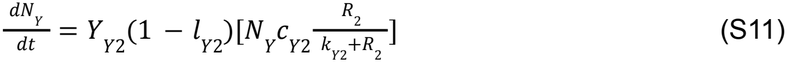

Replacing the bracket with (S3):

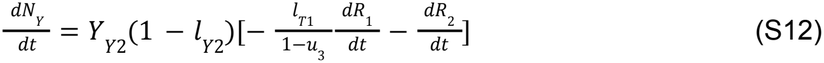

As in (S4), the *c*’s and *k*’s have cancelled: each biomass rate is a constant multiple of the resource rates.

#### Supplementary Text 3: Three-species models (TMY, TGY)

##### TMY model

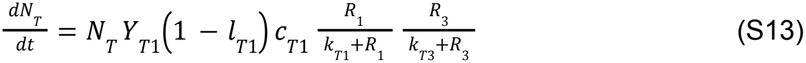

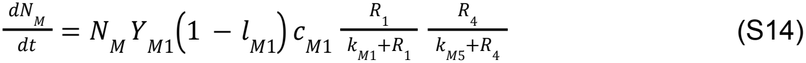

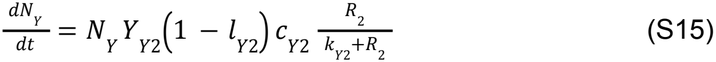

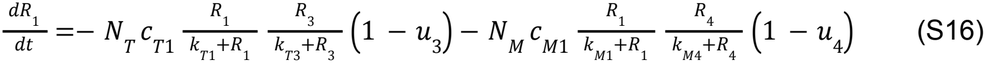

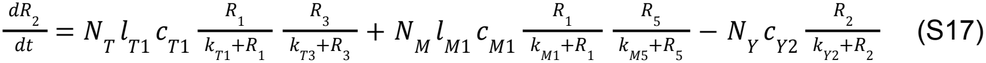

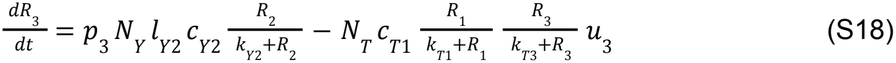

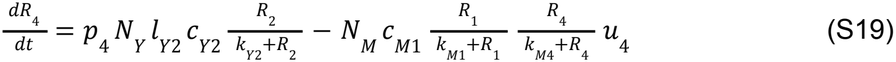

##### TGY model

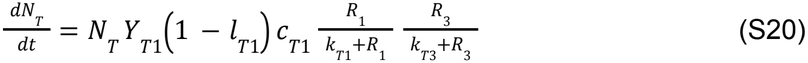

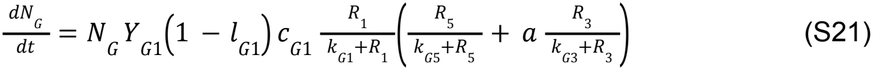

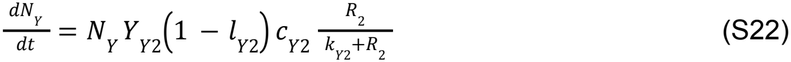

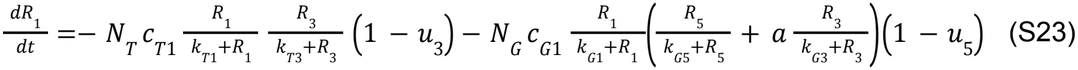

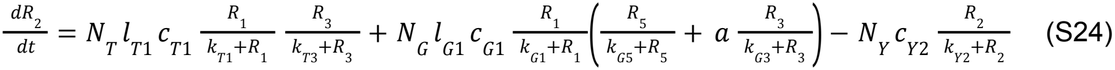

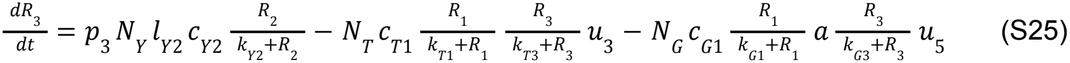

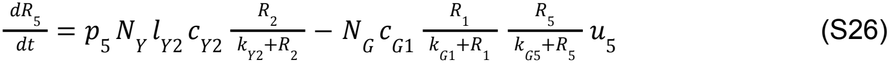

